# Development of Putative Isospecific Inhibitors for HDAC6 using Random Forest, QM-Polarized docking, Induced-fit docking, and Quantum mechanics

**DOI:** 10.1101/2020.08.10.243824

**Authors:** Ireoluwa Yinka Joel, Temidayo Olamide Adigun, Olukayode Olusola Bankole, Ahmeedah Ololade Ajibola, Emmanuel Bankole Ofeniforo, Faith Beyaan Auta, Ugochukwu Okechukwu Ozojiofor, Ifelolu Adeseye Remi-Esan, Aminat Ifeoluwa Akande

## Abstract

Histone deacetylases have been recognized as a potential target for epigenetic aberrance reversal in the various strategies for cancer therapy, with HDAC6 implicated in various forms of tumor growth and cancers. Diverse inhibitors of HDAC6 has been developed, however, there is still the challenge of iso-specificity and toxicity. In this study, we trained a Random forest model on all HDAC6 inhibitors curated in the ChEMBL database (3,742). Upon rigorous validations the model had an 85% balanced accuracy and was used to screen the SCUBIDOO database; 7785 hit compounds resulted and were docked into HDAC6 CD2 active-site. The top two compounds having a benzimidazole moiety as its zinc-binding group had a binding affinity of −78.56kcal/mol and −78.21kcal/mol respectively. The compounds were subjected to exhaustive docking protocols (Qm-polarized docking and Induced-Fit docking) in other to elucidate a binding hypothesis and accurate binding affinity. Upon optimization, the compounds showed improved binding affinity (−81.42kcal/mol), putative specificity for HDAC6, and good ADMET properties. We have therefore developed a reliable model to screen for HDAC6 inhibitors and suggested a series of benzimidazole based inhibitors showing high binding affinity and putative specificity for HDAC6.

## 1.0. Introduction

Cancer is a complex disease showing multifaceted and multifactorial presentation, with a high rate of morbidities and mortalities second only to cardiovascular disease.^[1]^ There is growing clarity on the significant effect of epigenetic alterations in cancer initiation and development. The post-translational modifications involve reversible acetylation and deacetylation of histone^[2][3]^ by adding or removing acetyl groups to or from specific lysine residues respectively by histone acetylases and deacetylases enzymes at the histone tails.^[4][5][6][7]^ This increases or decreases ionic interaction respectively between the positively charged histones and negatively charged DNA to yield more or less compact chromatin structure,^[8]^ thereby regulating gene expression and the subsequent accessibility of cell growth transcription machinery as well as RNA polymerase enzyme accordingly for gene transcription repression or enhancement and microtubule stability regulation in response to both intrinsic and extrinsic stimuli.^[9][8]^

The DNA histone deacetylation by Histone deacetylases results in tighter DNA-histone core wrapping, thereby leading to condensation of chromatin with its resultant gene expression silencing^[10]^ as well as gene transcription factors repression.^[11]^ This explains why over-expression of HDACs, resulting in abnormal levels of deacetylated histones and its associated normal gene transcription inhibition, have been linked to cancer cell proliferation,^[12][13]^ as different HDACs are over-expressed in many cancer cell lines and tissues, thereby indicating their crucial role in tumor development.^[14][10]^

The human histone deacetylase 6 belongs to Class IIb of the family of zinc-dependent hydrolases in the eighteen currently identified isoforms of histone deacetylase enzymes based on their sequence homology, sub-cellular distribution and catalytic activity.^[13][1]^ It has two catalytic domains including a ubiquitin-binding domain at the C-terminal region and a dynein-binding domain.^[15][16]^ Histone deacetylase 6 has been shown to have a wide range of abnormal differential expressions in solid tumors and hematological malignancies including prostate cancer, lung cancer, liver cancer, ovarian cancer, renal cancer, thyroid cancer, breast cancer, urothelial cancers, gliomas, lymphomas, etc.^[17][18][19][20]^, thereby rendering it a viable target for cancer therapy.^[21]^

There are currently many HDAC inhibitors or pan-inhibitors that have shown certain anti-tumor activity under experimental and clinical trials,^[22][23]^ however non-specificity, low bioavailability and potential for drug-drug interactions (through cytochrome P450 inhibition) has been a major challenge.^[1]^ The pharmacophoric features of HDAC inhibitors consist of a capping group (surface recognition moiety), a hydrophobic spacer (the linker), and a zinc-binding group (ZBG).^[24][25]^ The current ZBG ranges from hydroxamic acid, 2-aminoanilide, electrophilic ketones, to short-chain fatty acids.^[26]^ Bolden *et al*. (2006), Guo *et al*. (2012) and Lobera *et al*. (2018) suggested that changes in the cap group, the linker or the ZBG can individually provide selectivity for specific HDAC isoform inhibition^[27][28]^ as there is widespread speculation that development of isotype-selective inhibitors may contribute to therapeutic index improvement over non-selective ones.^[13]^

In this study, we aimed to identify, optimize potential HDAC6 specific inhibitors, and determine the mechanism of inhibition and specificity. The workflow for this study includes:

- building a Random Forest Classifier (RFC) model which “learned” from all know HDAC6 inhibitors that have been deposited in the Chembl database from 2003 to date, and validated extensively;
- screening the SCUBIDOO database (a database of 21,000,000 compounds set up for fast and efficient lead identification) for hit compounds;
- subjecting the hit compounds to Induced-fit docking, QM-polarized docking, QM-MM geometric optimization, and Molecular dynamics (Normal mode analysis), in other to determine binding affinity, elucidate a binding hypothesis (with interest in the mechanism of specificity and inhibition) and stability of the protein-ligand complex;
- optimization of compounds (Bioisostere replacement protocols) for improved binding affinity and specificity;
- calculate electronic descriptors (using Quantum mechanics) to determine the mode of chemical reactivity (nucleophilic or electrophilic) and stability of the optimized compounds;
- predict ADMET properties.

## 2.0. Material and Methods

### 2.1. Hardware and software

All analysis was carried out using a Linux Ubuntu 18.04 distro system running on a 12 GB RAM, core i5, 4 Cores, 2.5GHz.

All python packages were run on Python 3.6 using Jupyter Lab 1.2.6. Python packages used include Scikit-learn (v0.22.2) for model training, Feature Selector for feature extractions and preparations, Pandas (v1.0.3) for data waggling, Matplotlib (v3.2.1) and Seaborn (v0.10.0) for data visualizations, Numpy (v1.18.1), SHAP and PDPbox for model interpretation, TPOT (v0.11.1) for AutoML analysis.

### 2.2. Data Extraction and Preparation

All inhibitors of HDAC6 present in the ChEMBL^[29]^ database were downloaded (3,744) and imported into a standalone MySQL database we created for analysis. All inhibitors with missing IC50 values were removed, thereafter inhibitor smiles and corresponding IC50 values were extracted into a CSV sheet.

Using the MOE descriptor calculator, 2D descriptors were calculated for the inhibitors. A python script implementing Feature selector module was written to remove inter-correlated descriptors (correlation threshold was set at 0.75). Using the same module, all descriptors that did not contribute to 0.95 cumulative importance were removed (feature selector uses XGBoost algorithm to determine the feature importance of the descriptors).

A python script was written to convert IC50 values to pIC50 values (pIC50 = 9-log10(IC50)) (All IC50 were in nM). The pIC50 values were converted to categorical values of active (1) and non-active (0). The minimum pIC50 (2: the equivalent of 10000000 IC50 value) and maximum pIC50 value (12: the equivalent of 0.002nM IC50 value) were considered before setting activity threshold; this is to ensure balanced classes: Activity threshold was therefore set to 7 pIC50 and above (the equivalent of 37nM IC50 value). The data-set was finally divided into training, test, and external validation set.

### 2.3. Model building

A python script was written which implemented a Tree-based pipeline optimizer (TPOT).^[30]^ TPOT is an AutoML python module that searches for the best machine learning algorithm for our modeling task. It uses a genetic algorithm to search different tree-based algorithms (and corresponding best hyperparameter values), feature processing and selection algorithm, and outputs the pipeline with the highest accuracy. The following TPOT parameters were set: Generation: 100, Population Size: 100, Cross-validation: 10, and Early stopping: True. All other parameters were left at default values.

Random forest classifier (RFC) algorithm was the best performing algorithm with the following hyper-parameters: bootstrap: True, criterion: Gini, maximum features: 0.3, minimum samples leaf: 13, minimum samples split: 13, and number of estimators: 100.

Using Sci-kit learn RFC was fitted on the dataset and evaluated using Sci-kit learn classification evaluation metrics^[31]^ the following metric was calculated to evaluate the performance of the RFC model after 10 fold cross-validation: Recall (sensitivity), specificity, Precision, F1, Accuracy, Balanced accuracy, Error rate, and confusion matrix as described by Hossin *et al.*^[32]^

In other to provide a suitable explanation for the RFC model we wrote a python script implementing SHAP and PDPbox module. SHAP plotted feature importance of the descriptors^[33]^, PDPbox plotted the partial dependence plot (PDP), and individual conditional exception (ICE) plot.^[34]^

### 2.4. Database Screening

We used the SCUBIDOO database a freely available database of 21 million virtual compounds.^[35]^ The screening was in four steps:

- Predicting the activity of the M sample (∼99,000 compounds) representation of SCUBIDOO with the RFC model
- Docking of predicted active compounds to HDAC6 active site and identification of part/fragment of the molecule making binding interaction of interest (chelating interaction)
- Searching the SUBIDOO database for compounds having the identified fragments
- Hits from the search are downloaded for further analysis

### 2.5. Structure-based drug discovery

#### 2.5.1. Molecular docking

The crystal structure of HDAC6 in complex with TSA in the catalytic domain (CD) 2 (PDB: 5WGI) was downloaded from PDB. The protein was prepared using the Schrödinger protein preparation wizard^[36]^; missing side chains and loops were filled with prime,^[37]^ waters beyond 5Å from het group was deleted and het states were generated using Epik ^[38]^ (pH 7.0 +/- 2.0); all other parameters were left at default values.

The downloaded SCUBIDOO compounds were prepared using the Schrödinger Ligprep module; force field minimization using OPLS2005,^[39]^ het states generated using Epik (pH 7.0 +/- 2.0) and metal-binding states were added. All other parameters were left at default values.

Using the receptor grid generation module of Schrödinger the Grid file (active site coordinates and constraints) for the protein was extracted. The compounds were thereafter docked using Schrödinger’s virtual screening workflow. Briefly, the workflow consists of filtering (drug-likeness criteria), ligand preparation (using Ligprep), docking (the docking phase involved using the three Glide docking protocols: HTVS, SP, XP) and postprocessing (rescoring) using prime molecular mechanics-generalized Born surface area (MM-GBSA)^[40]^. All parameters were left at default values. The docking protocol was validated by redocking the co-crystalized ligand; the RSMD value was thereafter calculated.

#### 2.5.2. Molecular Mechanics Generalized Born Surface Area

Molecular Mechanics Generalized Born Surface Area (MM-GBSA) evaluated the binding free energies (binding affinity) and minimized the docked protein-ligand complex.^[41]^ Using VSGB 2.0 implicit solvation model and OPLS-2005.^[42]^ The binding free energy was calculated using Eq.1

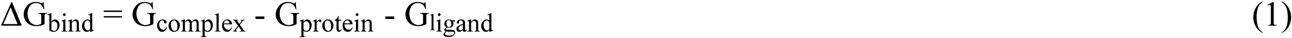

Where G_complex_, G_protein,_ and G_ligand_ represent the free binding energy of the protein-ligand complex, protein, and ligand respectively.

#### 2.5.3. Induced-fit docking

Schrodinger induced-fit docking module^[43][44]^ was used to perform Induce-fit docking. All parameters were left at default values. Briefly, the ligand is docked into the active site of the protein using Glide protocol (SP)[45] with the protein active site residues held rigid, next the protein side chains or backbone are refined using prime refinement module,^[37]^ finally, the ligand is redocked into the refined protein conformation and induced fit docking score is calculated to rank the protein-ligand complex.

#### 2.5.4. QM-polarized docking

QM-polarized docking (QPLD) was implemented using the Schrödinger QPLD module.^[46]^ Briefly, the ligands are docked into the active site of the protein using glide docking protocol^[45]^ and partial charges calculated by force field minimization, next the best-docked pose are selected and partial charges induced on the ligands by the active site residues are calculated using quantum mechanics this calculation is implemented using Schrödinger Qsite module,^[47]^ finally, the ligands are redocked using the QM derived partial charges and ranked using glide docking score.

#### 2.5.5. QM/MM optimization

Using the Schrodinger Q-site module QM/MM optimization was carried out on docked protein-ligand complexes. Briefly, the ligand and active site residue (side-chain and backbone) involved in the interactions are treated as the Quantum mechanics (QM) region while the other protein complex is treated as molecular mechanics (MM) region. The Qm calculation was done using Density functional theory (DFT) with Becke’s three-parameter exchange potential, Lee-Yang-Parr correlation functional (B3LYP) and basis set 631G^**^ level,^[41]^ while MM region was treated using OPLS2005; minimization was done using Truncated Newton, maximum cycle:1000 with convergence criterion set to Energy Gradient.^[41][47]^ All other parameters were left at default.

#### 2.5.6. Molecular dynamics

The stability of the docked protein-ligand complex investigated was using Normal mode analysis (NMA). The ease of deformability of the complex (eigenvalues), B-factor, variance, co-variance map, and the elastic network was calculated.^[48]^ Using iMOD server (http://imods.chaconlab.org) NMA calculations were implemented

#### 2.5.7. Bioisostere replacement

In other optimized the lead compounds Schrodinger Bioisostere replacement module was used. The module replaced certain functional groups with the ligands while retaining biological activity or improving activity.

### 2.6. Electronic descriptor calculations

Quantum mechanics (QM) theories were used to determine Electronic descriptors of the ligands. Using Schrodinger Jaguar single energy point module,^[49]^ the ligands were optimized geometrically using hybrid Density flow theory (DFT) with Becke’s three-parameter exchange potential, Lee-Yang-Parr correlation functional (B3LYP) and basis set 631G^**^ level. The following descriptors were calculated: Highest Occupied Molecular Orbital (HOMO) energy, Lower Unoccupied Molecular Orbital (LUMO) energy, and Molecular Electrostatic Potential (MESP). From the HOMO and LUMO calculations, the following descriptors were further extrapolated:

- HOMO-LUMO gap= E_LUMO_ − E_HOMO_;
- ionization energy (I) = −E_HOMO_;
- electron affinity (A) = −E_LUMO_;
- global hardness (η) = (− E_HOMO_+ E_LUMO_)/2
- chemical potential (µ) = (E_HOMO_+E_LUMO_)/2 can be determined.
- Parr *et al*. ^[50]^ proposed the global electrophilicity power of a ligand as ω = µ2/2η.

### 2.7. ADMET property prediction

Using the Discovery studio ADMET property prediction module ADMET descriptors were calculated for the compounds. MOE (Molecular Operating Environment 2015) descriptor module was used to determine Linpinkis’s drug-likeness and Opera lead-likeness of the compounds.

## 3.0. Results

### 3.1. Ligand-based drug discovery

#### 3.1.1. Data extraction and preprocessing

A total of 3742 HDAC6 inhibitors developed from 2003 to date with IC50 values ranging from 0.1 to 100000nM were downloaded from the CHEMBL database. The smiles of these inhibitors were extracted and a total of 206 2D descriptors were calculated using MOE software; the data was prepared and features extracted using a python script that implemented feature selector module (it removed inter-correlated, zero and low importance descriptors). Out of the 206 descriptors, 43 contributed to 0.95 cumulative importance and were selected for model building

#### 3.1.2. Tree-Based Pipeline Optimization Analysis

Different machine learning algorithms have been developed, each with its array of hyper-parameters that should be tuned in other to achieve accurate models suited for different datasets. This tuning however requires expertise, experience, coupled with trial and error which is generally challenging and time-consuming. In other to solve this problem Auto-machine learning (AutoML) modules have been developed, these modules exhaustively search different algorithms and hyper-parameters that best suit the modeling task at hand. One of these AutoML modules is Tree-Based Pipeline Optimization (TPOT).^[30]^

TPOT is an AutoML module that uses a genetic algorithm to search for tree-based algorithms, feature selection, and preprocessing algorithms with their corresponding hyper-parameter suitable for the provided data.^[50]^ TPOT classifier was therefore used to search for the best machine learning algorithm suited for our data (see methods). The analysis produced the Random Forest Algorithm with an accuracy of 0.83 after 100 generations of search, with hyper-parameter: criterion: Gini; maximum features: 0.3; minimum samples leaf: 13; minimum samples split: 13; number of estimators: 100.

#### 3.1.3. Random Forest Classifier Model Building and Evaluation

The dataset was divided into 3 (training, test, and external validation set) with a ratio of 60:20:20 respectively, with the above hyper-parameters we fitted the Random Forest Classifier (RFC) on the training data set and evaluated its performance on the test and external data-set. Confusion matrix (Figure S1) and other classification matrix were used to evaluate the model performance (Table 1). The training set had a precision: 0.89; recall: 0.94; accuracy: 0.90; and balanced accuracy: 0.90 (Table 1). The test set had a precision: 0.78; recall: 0.90; accuracy: 0.87; and balanced accuracy: 0.86 (Table 1). The external validation set had a precision: 0.87; recall: 0.91; accuracy: 0.87; and balanced accuracy: 0.86 (Table 1). A Roc curve (True-positive rate Vs. False-positive rate) of the RFC model performance on its training, test, and external validation set was plotted (Figure 1) and AUC scores determined (Table 1; Figure 1). The receiver operative curve (ROC) curve and Area under Curve (AUC) score evaluates how well a model picks a true positive ahead of a false positive in a pool of true positive and false positives.[51] The training set had an AUC of 0.91; test set an AUC of 0.83; external validation set an AUC of 0.86

**Table 1:** Model evaluation metrics.

| Metric | Training-set | Test-set | External-set |
| --- | --- | --- | --- |
| **Precision** | 0.89 | 0.78 | 0.87 |
| **Recall** | 0.94 | 0.90 | 0.91 |
| **Accuracy** | 0.90 | 0.82 | 0.87 |
| **F1** | 0.91 | 0.84 | 0.89 |
| **Balanced Accuracy** | 0.90 | 0.83 | 0.86 |
| **ROC-AUC** | 0.91 | 0.83 | 0.86 |

**Figure 1:**
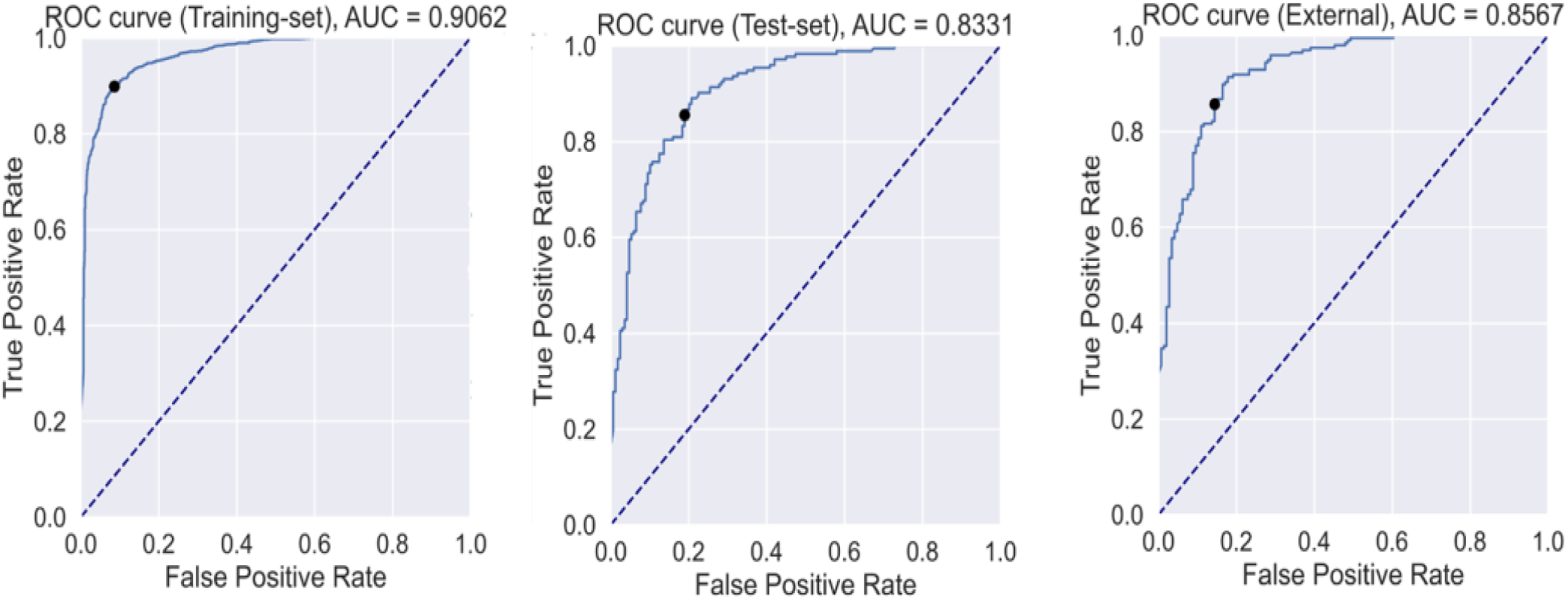
ROC-curve for Training, Test, and External validation data-set.

#### 3.1.4. Cross-Validation

Using Sci-Kit learn cross-val-predict function a 10 fold cross-validation was implemented on the datasets (Table 2). The training set had a precision: 0.83; recall: 0.39; accuracy: 0.83; and balanced accuracy: 0.83 (Table 2). The test set had a precision: 0.74; recall: 0.85; accuracy: 0.78; and balanced accuracy: 0.77 (Table 2). The external validation set had a precision: 0.79; recall: 0.90; accuracy: 0.80; and balanced accuracy: 0.78 (Table 2). The cross-validation Roc curve of the RFC model performance on its training, test, and external validation set was plotted (Figure S2) and AUC scores determined (Table 2). The Area under Curve (AUC) score was evaluated, the training set had an AUC: 0.83; test set: 0.77; external validation set: 0.77. Based on these results, we held the model to be a robust and reliable model for screening.

**Table 2:** Cross-validation Evaluation Metrics.

| **Metric** | **Training-set** | **Test-set** | **External-set** |
| --- | --- | --- | --- |
| **Precision** | 0.83 | 0.74 | 0.79 |
| **Recall** | 0.89 | 0.85 | 0.90 |
| **Accuracy** | 0.83 | 0.78 | 0.80 |
| **F1** | 0.86 | 0.78 | 0.83 |
| **Balanced Accuracy** | 0.83 | 0.77 | 0.78 |
| **ROC-AUC** | 0.83 | 0.77 | 0.77 |

#### 3.1.5. Model Interpretation

Understanding how a model is using inputted descriptors to make predictions about the dependent variable (biological activity) is not only important in optimizing the model but also can give insights on dataset improvement (i.e. descriptors to improve for better activity).^[34]^ Using SHAP and PDPbox python module we plotted feature importance plot, partial dependence plot (PDP), Individual conditional expectation (ICE), and partial dependence interaction plots (see methods). SHAP module plotted the feature importance plot for the RFC model and showed the importance in this order GCUT_SLOGP_0, PEOE_VSA-1, SMR_VSA2, pKA, GCUT_SLOGP_2, etc. (Figure 2). It should be noted however that the contribution of these descriptors is towards class probability. Therefore, figure 2 shows the importance and contribution towards the probability of both classes modeled (class 1: active (blue), class 0: non-active (red))

**Figure 2:**
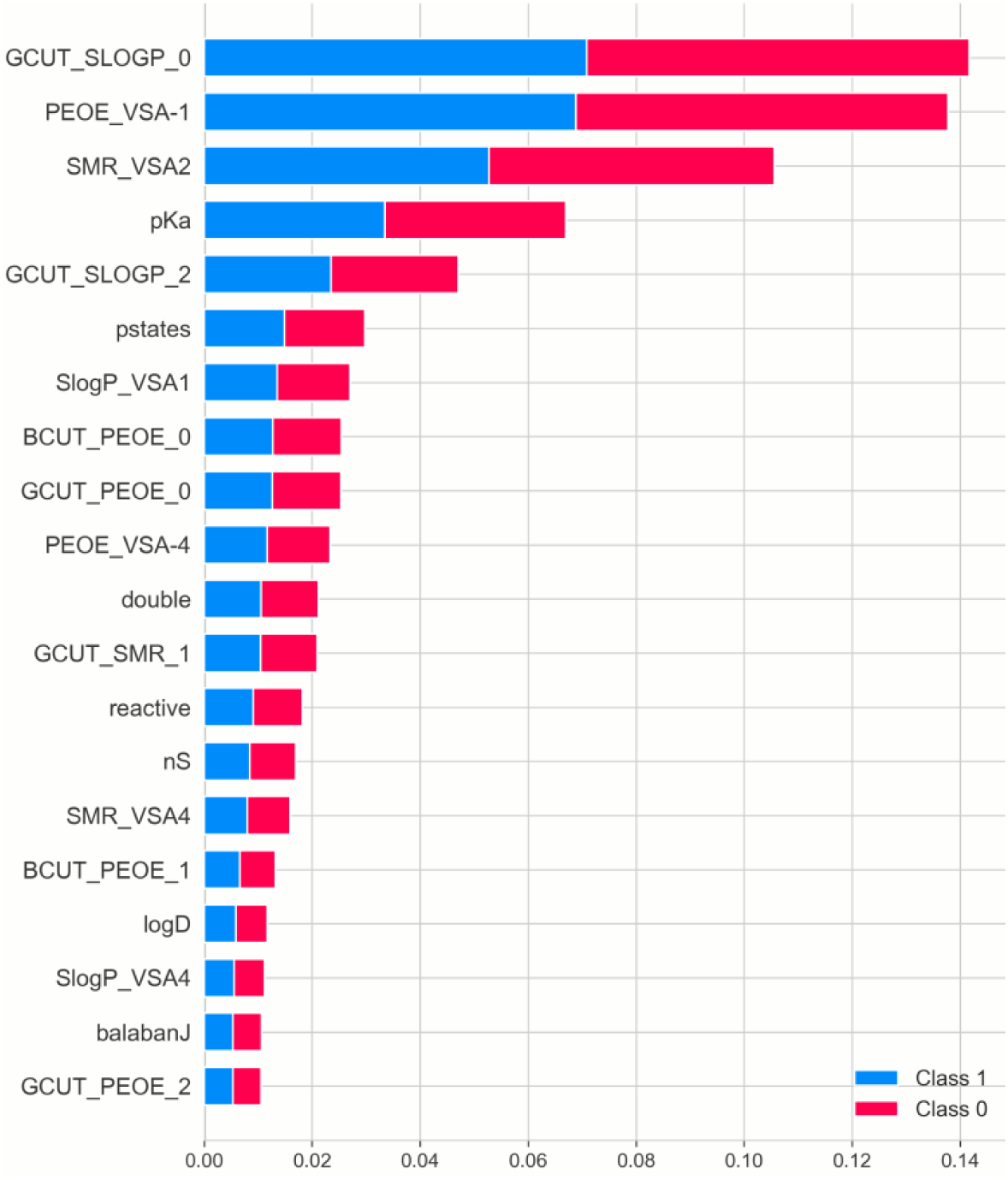
Random Forest Classifier feature importance.

The feature importance plot is one important plot for understanding machine learning model predictions however; it does not show how varying values of individual descriptors affect the model predictions. The partial dependence plot (PDP) solves this problem by showing the marginal effect of varying values of a descriptor on model prediction.^[52]^ The PDP showed that increasing values of PEOE_VSA-1, SMR_VSA2, and GCUT_SLOGP_2 increased RFC model active class probability predictions (Figure S3). This is, however, not true for GCUT_SLOGP_0 and pKA descriptors, increasing values of GCUT_SLOG_0 resulted in a negative contribution towards the RFC model active class probability contribution. pKa descriptors contributed positively only within a narrow range (−10 to 12) (Figure S3a[i], S3d[i]).

Partial dependence plot goes a step forward in providing insight into the ‘black-box’ of the RFC model however, the PDP plot is an average of all data points, this hides possible variations. Individual Conditional Expectation (ICE) plot, therefore, disintegrates the PDP plot into their corresponding data points showing possible trend variation.^[52]^ Using the PDPbox module we plotted ICE plots for the top five descriptors (Figure 5). The ICE plot suggests that although most of the data point followed the PDP plot trend some however deviated, for example, the PDP plot indicated that increasing values of GCUT_LOGP_0 resulted in negative contribution towards active class probability prediction but the ICE plot has however shown that this is not true for all data points (this also applies to pKA) (Figure S4).

Going a step further in trying to make the model prediction process as clear as possible we investigated the partial dependence interaction matrix. This matrix shows how various values of descriptors when considered together affect the active class prediction probability (this, therefore, reveals optimum values for active class prediction probability); the matrix considered two features/descriptors at a time. Overall GCUT_LOGP_0 optimum values for various interaction consideration was −1.17, pKa optimum value was 8.8, PEOE_VSA-1 optimum value was 121.93, SMR_VSA2 optimum value was 71.78, and GCUT_SLOGP_2 optimum value was 0.33 (Figure 3).

**Figure 3:**
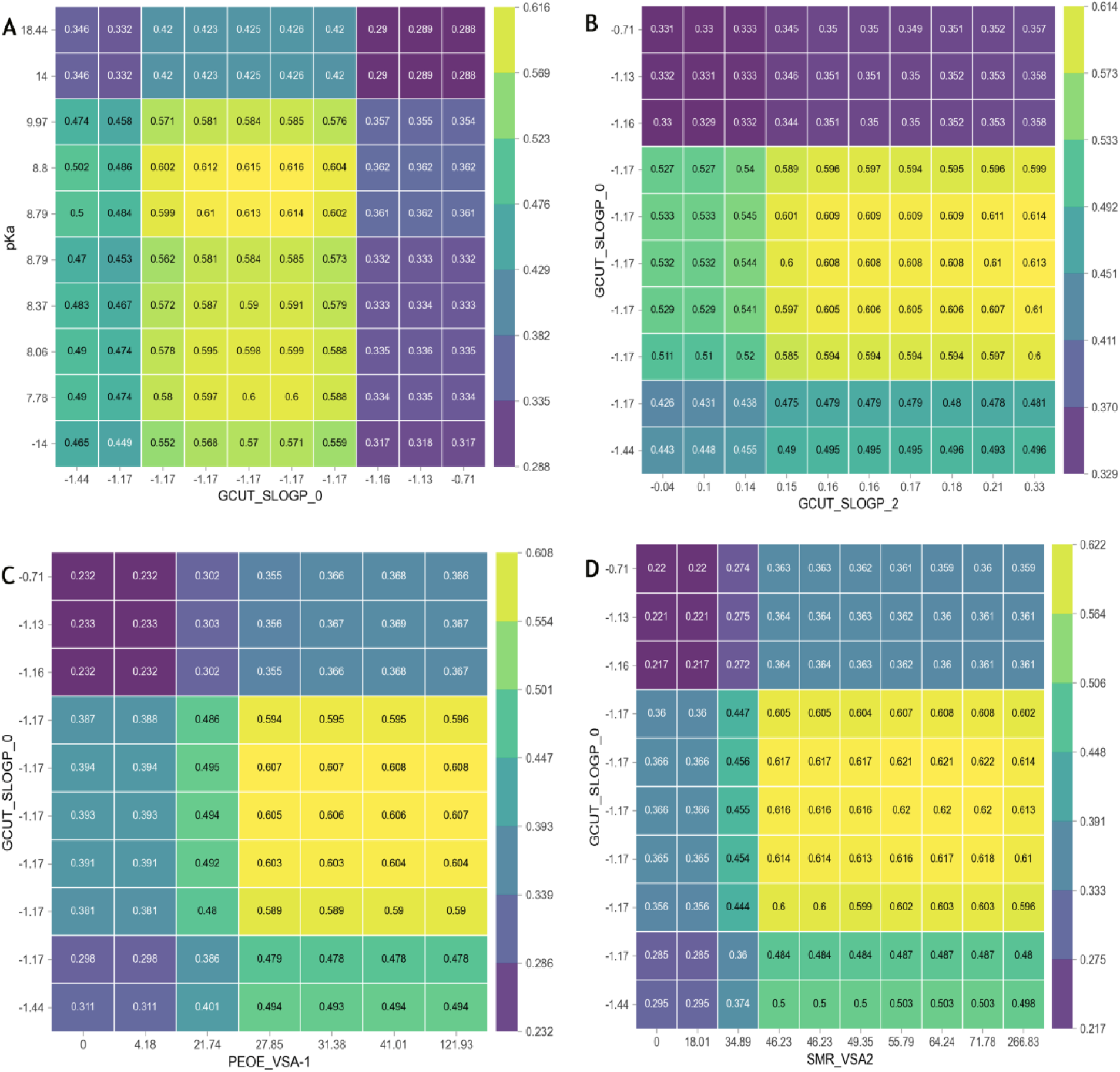
Partial dependence Interaction plot of. a) pKA and GCUT_SLOG_0 b) GCUT_SLOGP_0 and GCUT_SLOG_2 c) GCUT_SLOGP_0 and PEOE_VSA-1 d) GCUT_SLOGP_0 and SMR_VSA2

The model interpretation analysis has shown that GCUT_SLOGP_0 is the most important descriptor for our RFC model with an optimum value of −1.17 and that increasing values of this descriptor resulted in a negative contribution towards active class prediction. This, therefore, suggests that an increase in GCUT_SLOGP_0 values might result in a reduction in the biological activity of HDAC6 inhibitors. pKa was the third important descriptor and followed a similar trend with GCUT_SLOGP_0 with an optimum value of 8.8, increasing biological activity through pKa is however within a narrow range after which biological activities drop. Other descriptors however increased biological activity as their values increased. It should be noted that this analysis is suggestive and specific only to this model, this, therefore, means that other machine learning algorithms might result in deferent results hence experimental verification is still needed (Table S1 describes the descriptors used in developing the model).

#### 3.1.6. Database Screening

SUBIDOO a database with 21,000,000 virtual compounds/products^[35]^ was selected for screening, majorly for two reasons: the database has been set up in such a way as to reduce the computational cost of screening (see methods) and all compounds in the database come with detailed information on its synthesis and possible side reaction.

The RFC model was used to screen the M sample of the SCUBIDOO database for active compounds. Of the 99,977 compounds, only 167 compounds were classified as active; the 167 compounds were docked into the active site of HDAC6. The compound was filtered using the following criteria: docking score: > −8.0kcal/mol and interactions with catalytic ZN^2+^ ion. Two compounds were selected having met these criteria (docking score: −9.216 and −9.034kcal/mol respectively; formation of salt bridges with catalytic Zn^2+^ ion (Figure 4)). The moiety of these compounds making interaction with the ZN^2+^ was the hydropyrimidine functional group and thiadiazole functional group respectively. The SCUBIDOO database was, therefore, searched for compounds containing these functional groups and a total of 7785 compounds resulted from the search (Figure S4 shows the workflow for screening the database).

**Figure 4:**
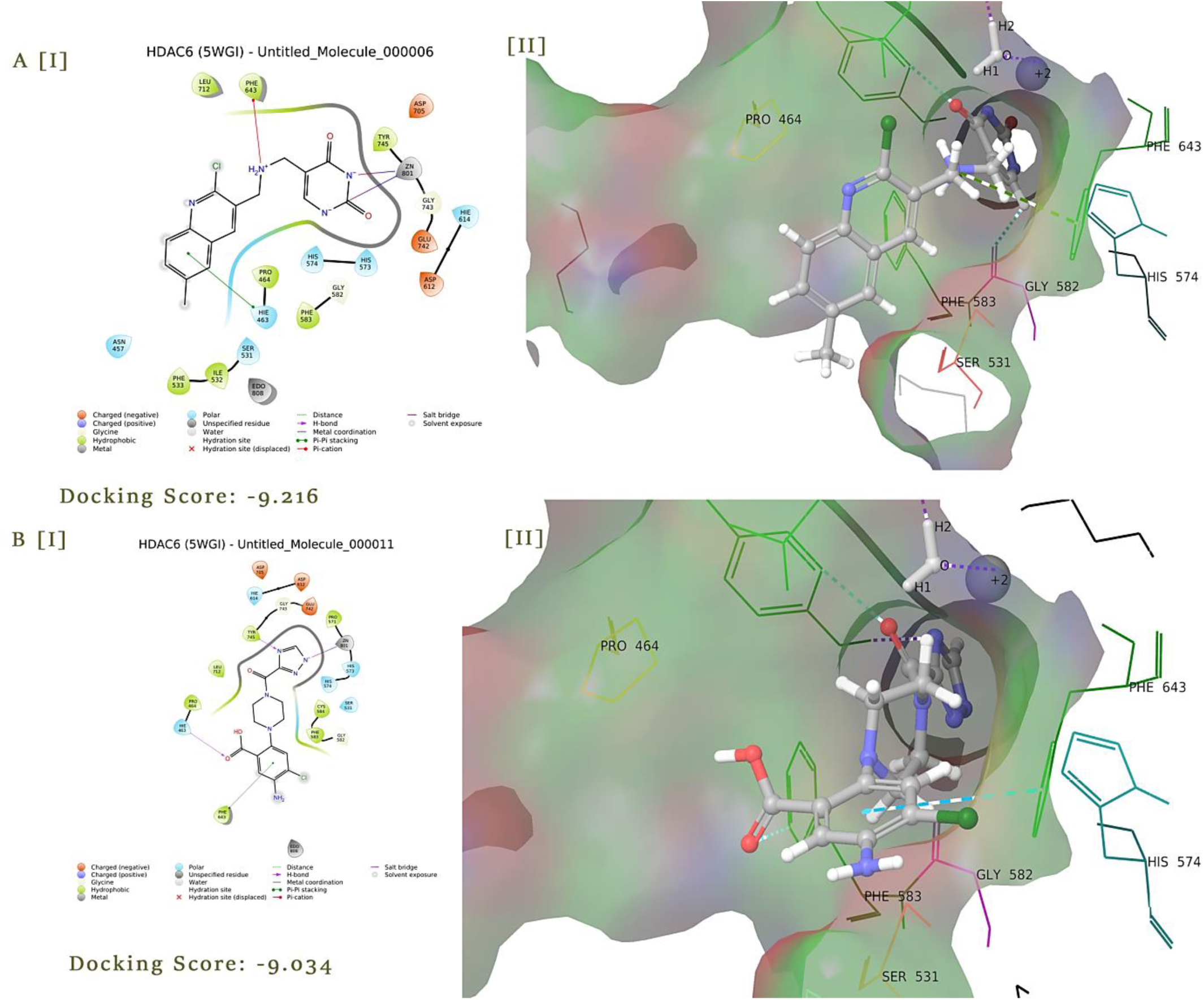
2D and 3D interaction of top best two M sample compounds docked in HDAC6 (CD2) active site: a) compound 6 b) compound 11

### 3.2. Structure-based drug discovery

#### 3.2.1. High throughput Virtual Screening

Using Schrodinger virtual screening workflow (see methods), the 7785 compounds downloaded from the previous section were docked into the catalytic domain 2 (CD2) of HDAC6. 24 compounds were returned with binding affinity ranging from −78 to −28kcal/mol (Table 2). Compounds 14660440 and 10651887 were the top compounds with the binding affinity of − 78.56kcal/mol and −78.21kcal/mol and were thus selected for this study (Figure 5). The docking protocol was validated by redocking the co-crystalized ligand TSA. The redocked pose was superimposed on the crystallized structure; an RSMD score of 1.60 was obtained (Figure S6).

**Figure 5:**
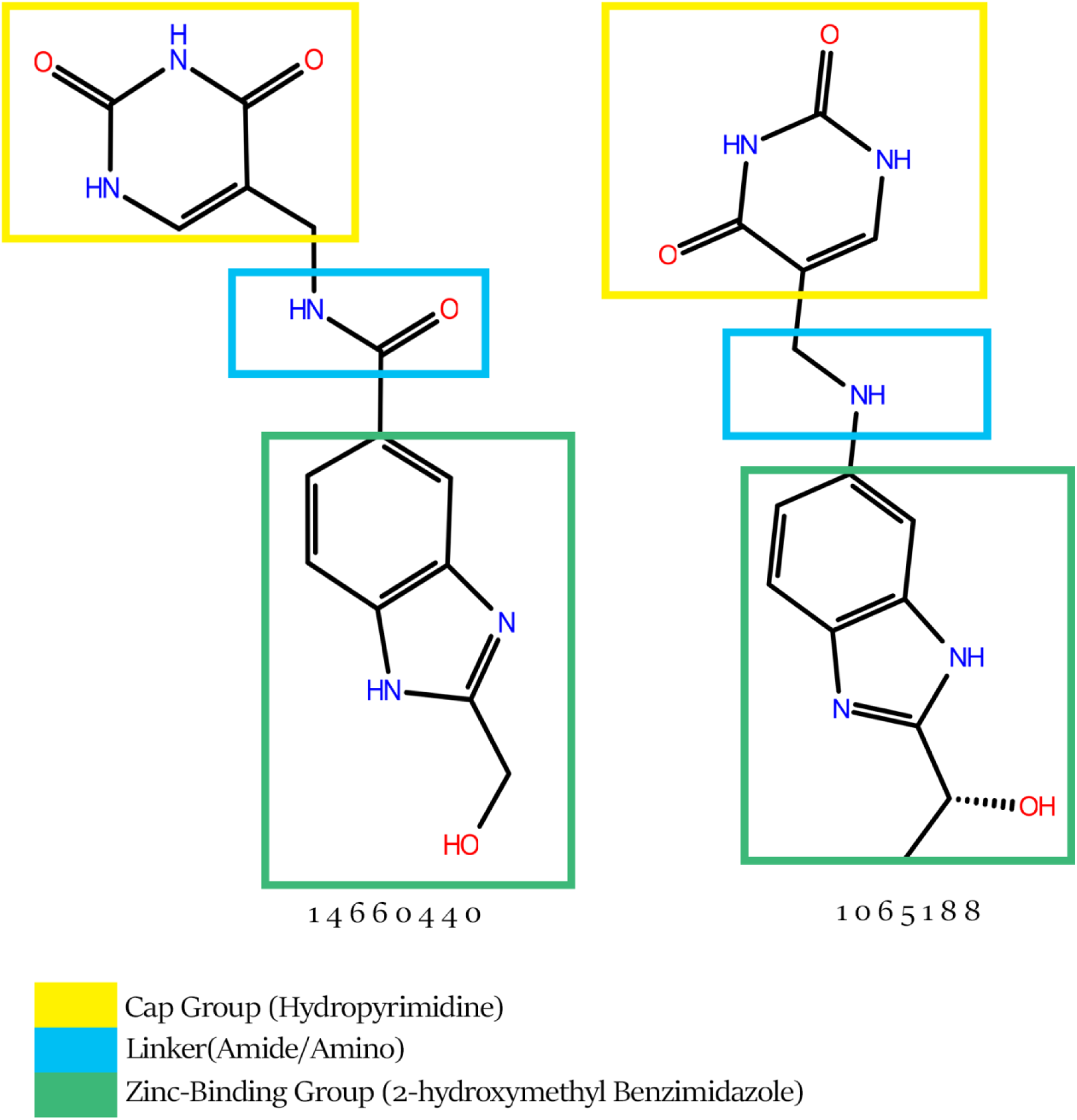
Compounds 14660440 and 1065188.

An inhibitor of HDAC6 which has shown very high selectivity (CAY10603)[52] and inhibitory activity for HDAC6 (IC50: 2 pM) was selected as the control for this study. When docked it had a binding affinity of −90.70kcal/mol (Table 3; Figure 6).

**Table 3:** Molecular docking, Induced fit docking, and Binding affinity.

| **Compound ID** | **XP-Docking score**<br>**(kcal/mol)** | **$\Delta G_{XP}$**<br>**(kcal/mol)** |
| --- | --- | --- |
| **CAY 10603** | -8.494 | -90.70 |
| **14660440** | -10.891 | -78.56 |
| **10651887** | -9.778 | -78.21 |
| **20558972** | -9.408 | -69.83 |
| **5899732** | -9.519 | -69.75 |
| **2559755** | -9.493 | -68.47 |
| **10640503** | -9.493 | -68.47 |
| **8964976** | -10.251 | -66.42 |
| **5886114** | -10.162 | -62.07 |
| **14742603** | -9.593 | -61.29 |
| **5127713** | -9.795 | -59.05 |
| **19366888** | -9.254 | -57.35 |
| **3646127** | -9.318 | -56.07 |
| **1481504** | -9.668 | -55.84 |
| **686879** | -9.429 | -50.26 |
| **18504893** | -9.63 | -48.99 |
| **10688493** | -9.118 | -46.05 |
| **5814288** | -9.664 | -45.63 |
| **11634999** | -10.176 | -43.57 |
| **3284098** | -9.191 | -43.43 |
| **5127628** | -9.604 | -67.86 |
| **7227152** | -10.053 | -27.44 |
| **3789238** | -9.544 | -29.39 |

**Figure 6:**
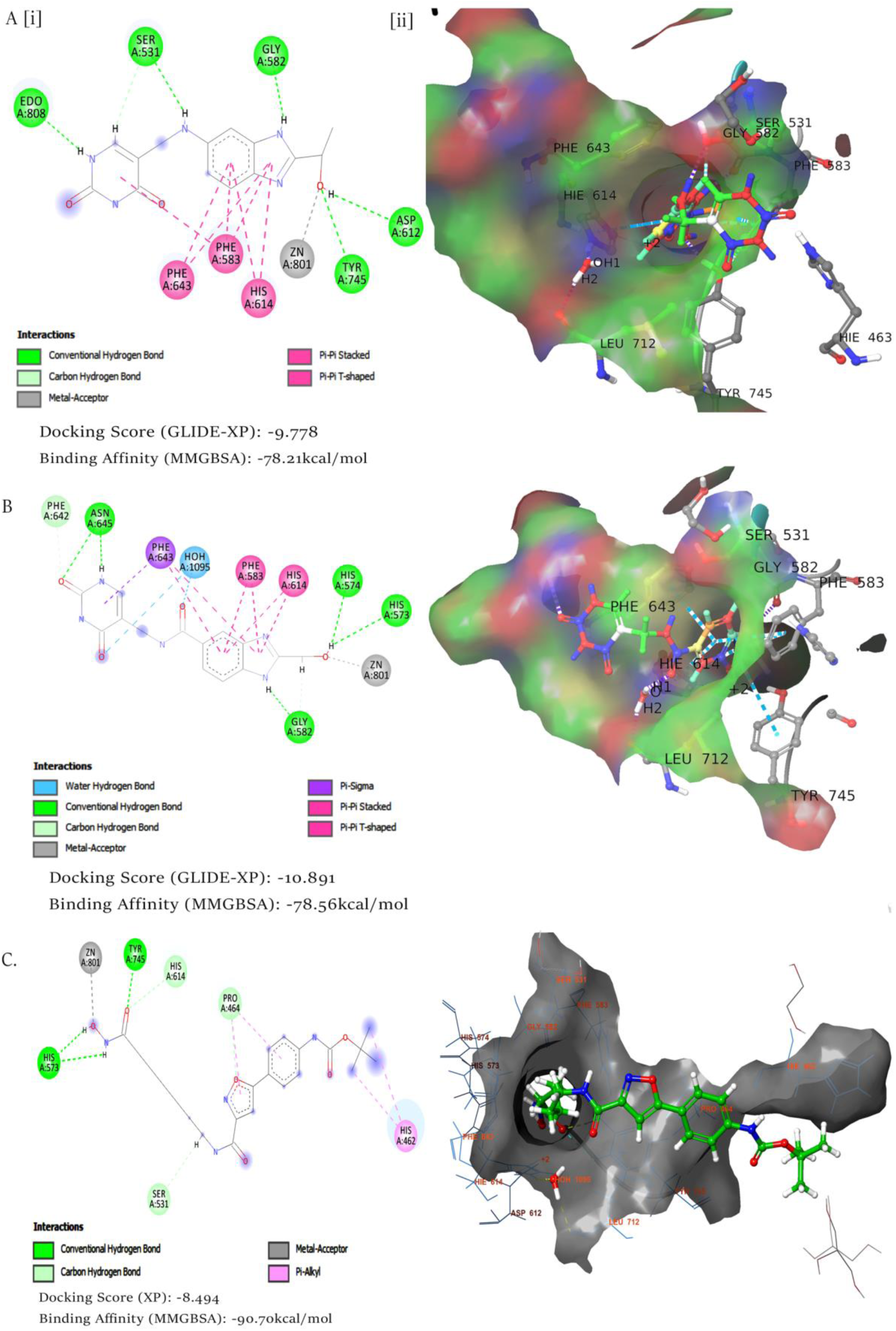
Molecular docking of compound a) 14660440 b) 10651887 c) CAY10603: i) 2Dinteractions ii) 3D interactions

Compound 14660440 formed a metal acceptor bond with the catalytic ZN^2+^ ion and Hydrogen Bond (H-bond) formation with HIS 573, HIS 574 using its hydroxymethyl R group. H-bond with catalytic water molecule via its amide linker and hydropyrimidine cap (Figure 6); HIS614 formed hydrophobic interactions (π-π bond) with the benzimidazole group. PHE 583, PHE643 also formed hydrophobic bonds (π-π, π-sigma) with the ligand; the hydropyrimidine cap formed H-bonds with PHE642 and ASN645 (Figure 6). Compound 10651887 formed metal acceptor bond with ZN^2+^ ion, H-bond with catalytic TRY745, H-bond with ASP612 and π-π interaction with HIS614 (both coordinators of the ZN^2+^ ion), π-π interactions with PHE583 and PHE643, H-bond with GLY582. The hydropyrimidine cap and amine linker formed H-bond with SER531 (Figure 6).

When comparing the binding poses of compound 14660440 and 10651887 with the CAY10603 (standard), the compounds had similar catalytic residue interactions. However, specificity differences are observed; interactions with PRO464 (H-bond, π-alkyl) and HIS462 (π-alkyl) (Figure 6). These two residues (PRO464 and HIS462) along with SER531 are specific only for HDAC6^[54]^; interactions with these residues, therefore, suggest high specificity for HDAC6 which is consistent with experimental data which has shown CAY10603 to be highly specific for HDAC6^[3]^ and also suggest the reason for its high binding affinity (−90.70kcal/mol) observed.

Having identified these compounds, we decided to further subject the compounds to more rigorous and accurate docking protocols. This was in a bid to elucidate a binding hypothesis (i.e. mechanism of inhibition and specificity). We, therefore, subjected the compounds (14660440 and 10651887) to more accurate and exhaustive docking protocols (Induced fit docking and QM-polarized docking) and geometrically optimize the docked complex using QM/MM optimization while comparing with the standard.

#### 3.2.2. QM-polarized docking

Accurate charges have been shown to improve docking accuracy,^[53]^ most docking protocols consider the charges on the ligand via force field minimization, however, charges induced on the ligand within the protein active site is not considered. QM-polarized docking (QPLD) however tries to improve docked ligand binding conformation by considering plausible charge polarization that might occur due to the active site residues.^[54]^ Using Schrödinger’s QPLD module (see methods) QPLD was implemented on compound 14660440, 10651887, and CAY10603 to gain insight into the binding interactions of these compounds and calculate their corresponding binding affinity. The results showed CAY10603 to have the highest binding affinity of −98.55kcal/mol. Compound 10651997: −75.83kcal/mol and compound 14660440: − 70.80kcal/mol (Table 4). New H-bond was observed between ASP612 and compound 10651887 other bonds remained when compared with the molecular docking. Compound 14660440 formed a new H-bond with SER531 however, this bond had a bond length of 2.78Å which was rather weak when compared to 2.49Å (Table 7) of compound 10651887 (Figure 7). CAY10603 formed H-bond with the catalytic water molecule, pi-alkyl bonds with LEU712, and alkyl bonds with LEU466 (Figure 7). The results showed that polarized charges induced by the active site residues on the ligands cause a reduction in the binding affinity of compound 10651887 and 14660440 for HDAC6.

**Table 4:** QPLD docking score and binding affinity.

| Compound ID | XP docking score(kcal/mol) | QPLD docking score (kcal/mol) | $\Delta G_{QPLD}$ (kcal/mol) |
| --- | --- | --- | --- |
| **CAY 10603 (Standard)** | -8.494 | -8.810 | -98.55 |
| **10651887** | -9.778 | -10.450 | -75.83 |
| **14660440** | -10.891 | -10.422 | -70.80 |

**Figure 7:**
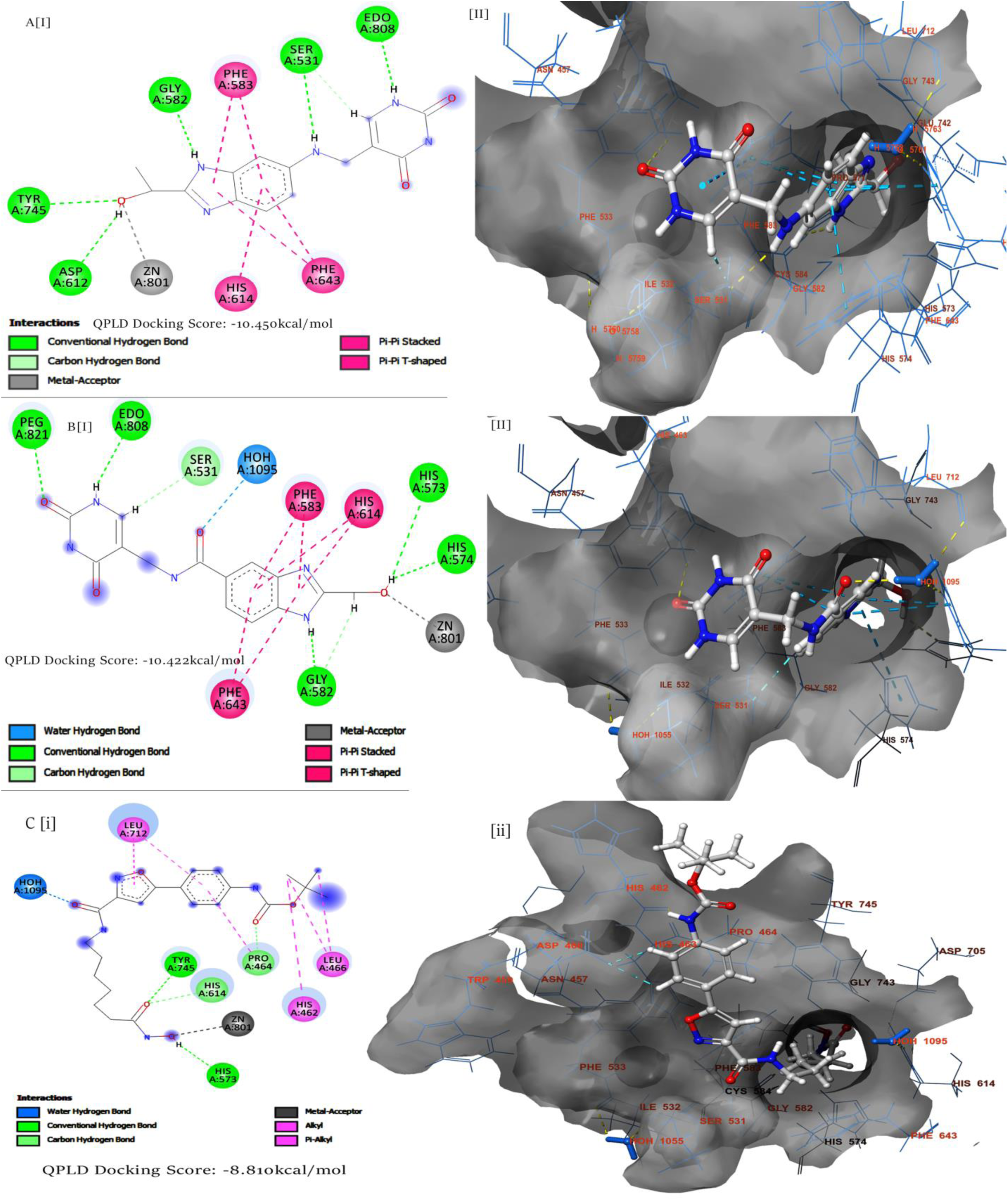
QM-polarized ligand docking of a) compound 10651887 b) compound 14660440 C) CAY10603: [I] 2D interactions [II] 3D interactions

#### 3.2.3. Induced fit docking

Apart from accurate charges, conformational changes in the active site as a result of ligand binding needs to be considered before a reliable binding hypothesis can be elucidated. Induced fit docking (IFD) considers both the ligand and active site residues flexibility in its calculations, this, increases the accuracy of predicted binding interactions and binding affinity.^[55][56]^ We performed IFD (see methods) on compounds 14660440, 1065188, and CAY10603 and calculated binding affinity.

Compound 10651887 had the highest IFD score of −847.34; two new bonds were observed: H-bond with HIS 574 and HIS573, π-Donor H-bond with PRO464, and Carbon H-bond with HIS463 were also observed. However, H-bond formed previously with Asp612 was lost, all other bonds were retained (Figure 8). Binding affinity however increased from −78.21kcal/mol (XP) to −98.29kcal/mol (Table 5) becoming the ligand with the highest binding affinity. Compound 14660440 had an IFD score of −847.20; of note is the formation of a new H-bound was formed with TYR745 all other bonds were retained (Figure 8). The binding affinity also increased to −89.63kcal/mol (Table 5). CAY10603 formed new H-bonds with GLY743 and GLY582, π-sigma bond with PHE643, π-π interaction .with PHE583 and alkyl bonds with LEU466; H-bonds with TYR745, HIS573, and HIS614 were lost (Figure 8). The binding affinity of CAY10603 was however reduced from −90.70kcal/mol to −85.74kcal/mol (Table 5).

**Table 5:** IFD score and binding affinity.

| Compound ID | IFD score | $\Delta G_{\text{IFD}}$ (kcal/mol) |
| --- | --- | --- |
| **CAY 10603** | -839.57 | -85.74 |
| **14660440** | -847.20 | -89.63 |
| **10651887** | -847.34 | -98.29 |

**Figure 8:**
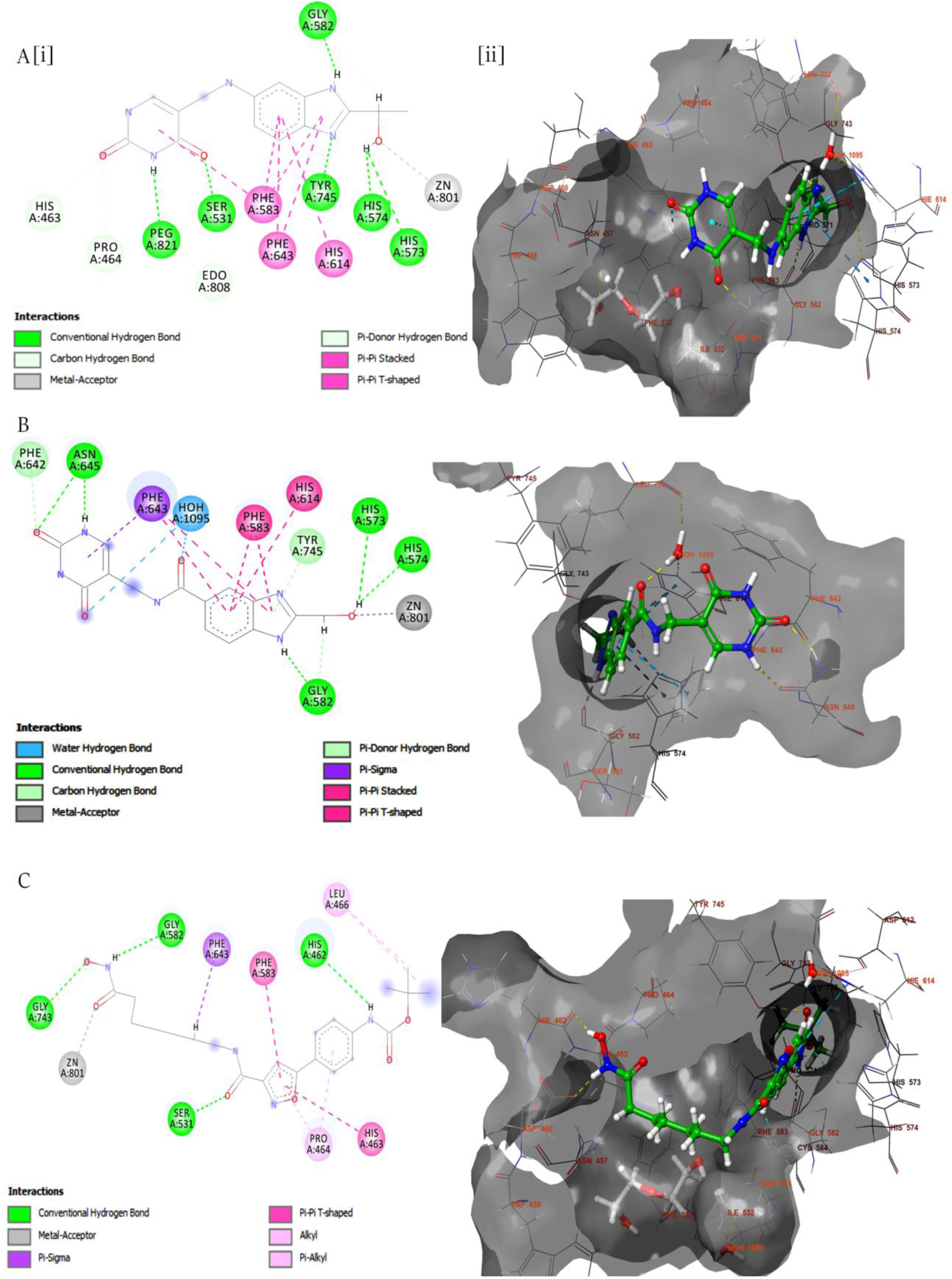
Induced fit docking of compound a) 10651887 b) 14660440 C) CAY10603: i) 2D interaction ii) 3D interaction

#### 3.2.4. QM/MM optimization

The induced-fit protein-ligand complex was optimized using QM/MM geometric optimization in other to validate the binding interactions observed. In QM/MM optimization protocol, we treated the ligand and interacting side chains as the QM region while the whole protein was treated as the MM region (see methods).

The QM/MM optimized pose was compared with the QPLD and IFD pose of the compounds. Compound 1466044 lost H-bonds formed with HIS 574 in the previous docking (IFD and QPLD) and reformed H-bonds with TYR745 which was missing in QPLD pose (Figure 9). Examining the bond length of the H-bond formed with HIS 573 and TYR 745 we observed stronger H-bonds upon optimization (Table 7); compound 10651997 retained H-bonds with HIS573, HIS574 which was absent in the QPLD pose (Figure 9). A new H-bond was formed with PRO464 which was not reported in any of the previous docking poses (IFD and QPLD).

**Figure 9:**
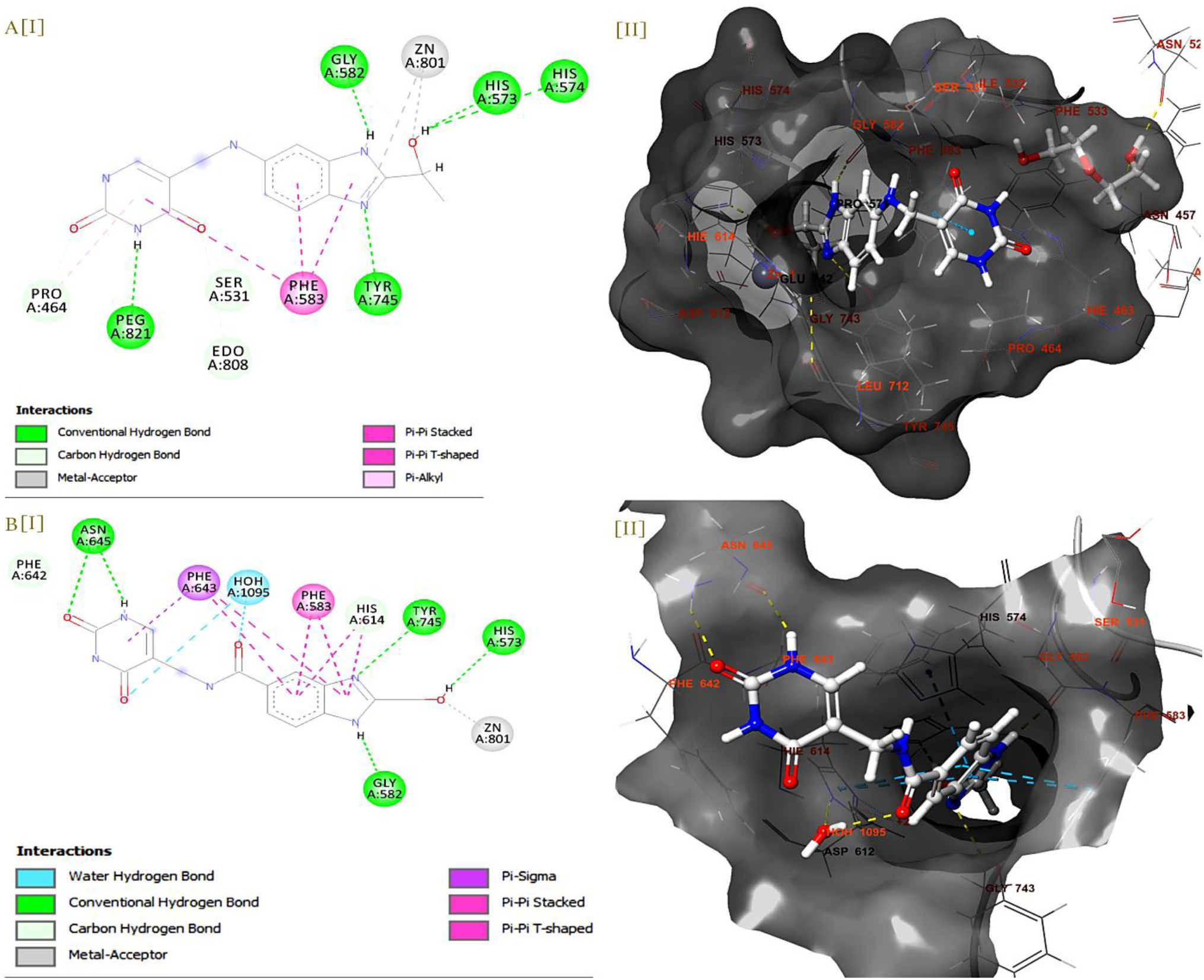
QM/MM optimization of protein-ligand docked complex: a) 10651997 b) 14660440

The H-bond strength with HIS573 improved upon optimization, however, H-bond with HIS574 was not as stable as previous pose (increase in bond length) (Table 7). H-bond formed with SER531 was retained but not as strong as the previous docking pose (Table 7). H-bond with TYR745 improved when compared with the QPLD pose. The binding affinity of compound 10651887 was −79.62kcal/mol and compound 14660440 was −77.44kcal/mol (Table 6). Over the for binding affinities calculated (Molecular docking, induced-fit, Qm-polarized docking and QM/MM optimization) compound 10651887 had the highest binding affinity when compared to compound 14660440 (table 6)

**Table 6:** Binding affinity of all docking protocols.

| **Compound ID** | **$\Delta G_{XP}$**<br>**(kcal/mol)** | **$\Delta G_{IFD}$ (kcal/mol)** | **$\Delta G_{QPLD}$**<br>**(kcal/mol)** | **$\Delta G_{QM/MM}$**<br>**(kcal/mol)** |
| --- | --- | --- | --- | --- |
| **CAY 10603** | -90.70 | -85.74 | -98.55 | - |
| **10651887** | -78.21 | -98.29 | -75.83 | -79.62 |
| **14660440** | -78.56 | -89.63 | -70.80 | -77.44 |

**Table 7:**
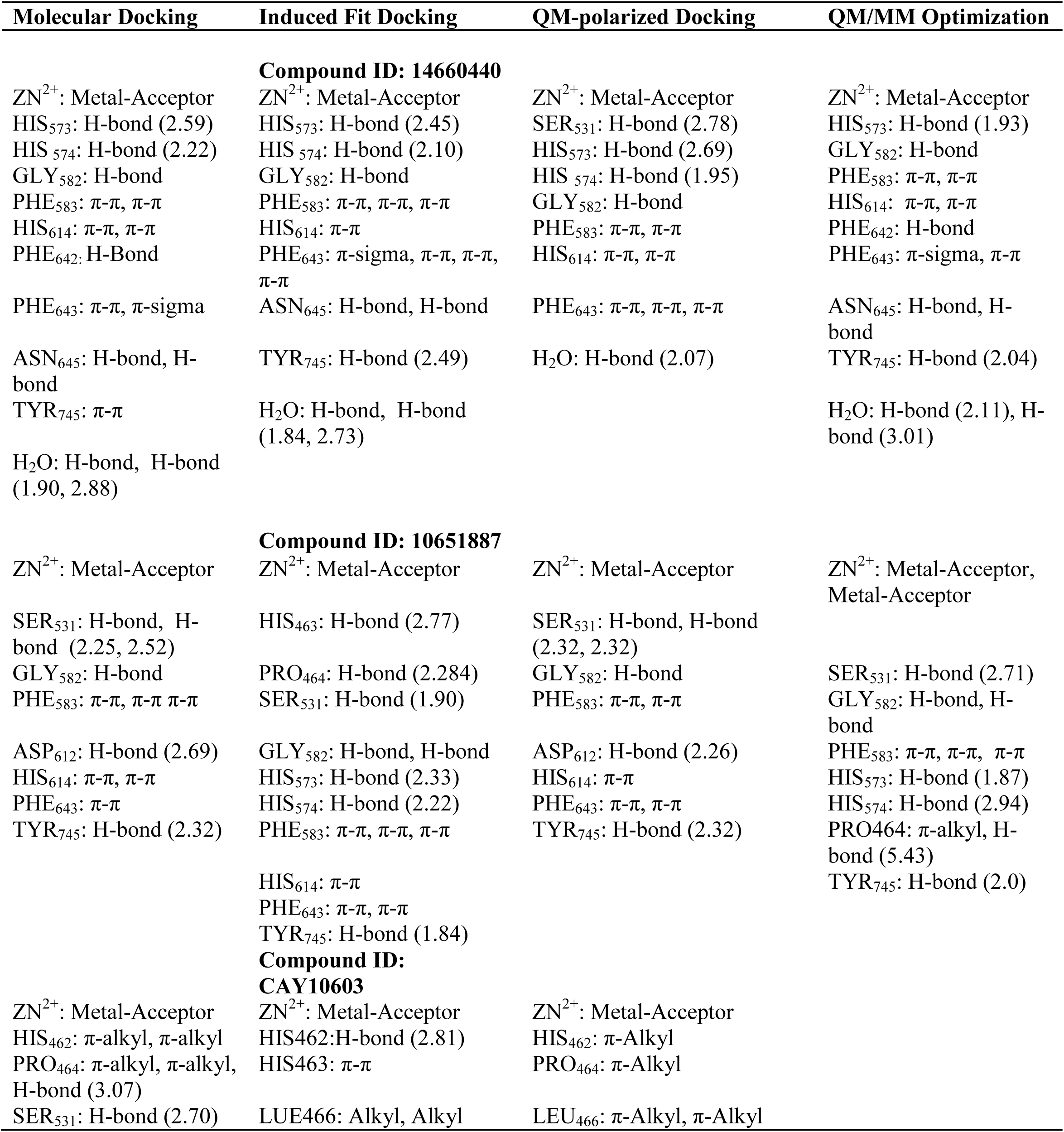
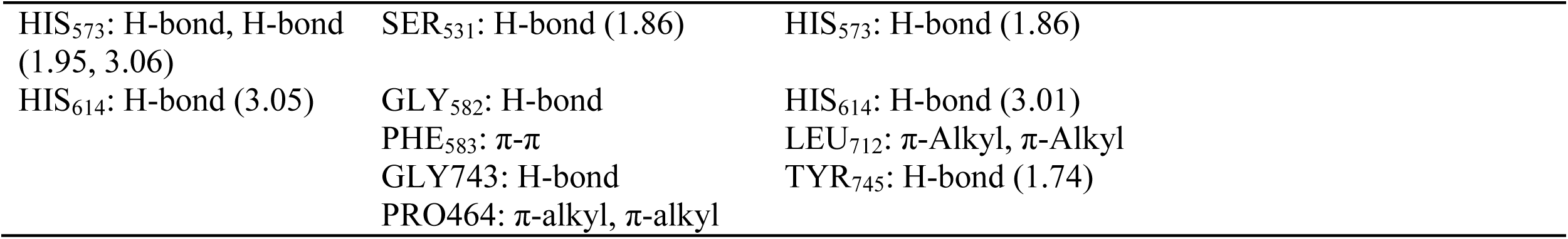
Summary of binding interaction of compound 14660440, 10651887, and CAY10603 over various docking protocol and optimization.

### 3.3. Molecular Dynamics

Using Normal mode analysis (NMA) the stability of the induce-fit docked complex, and ease of protein structure deformability was investigated. Using the elastic network model (ENM)[57] method, the degree of deformability of each residue (figure 10i), B-factor obtained from PDB field and NMA mobility data (figure 10ii), eigenvalues representing ease of deforment (figure 10iii), covariance map (figure S7), and elastic network (figure S8) for the docked complex was calculated.

**Figure 10:**
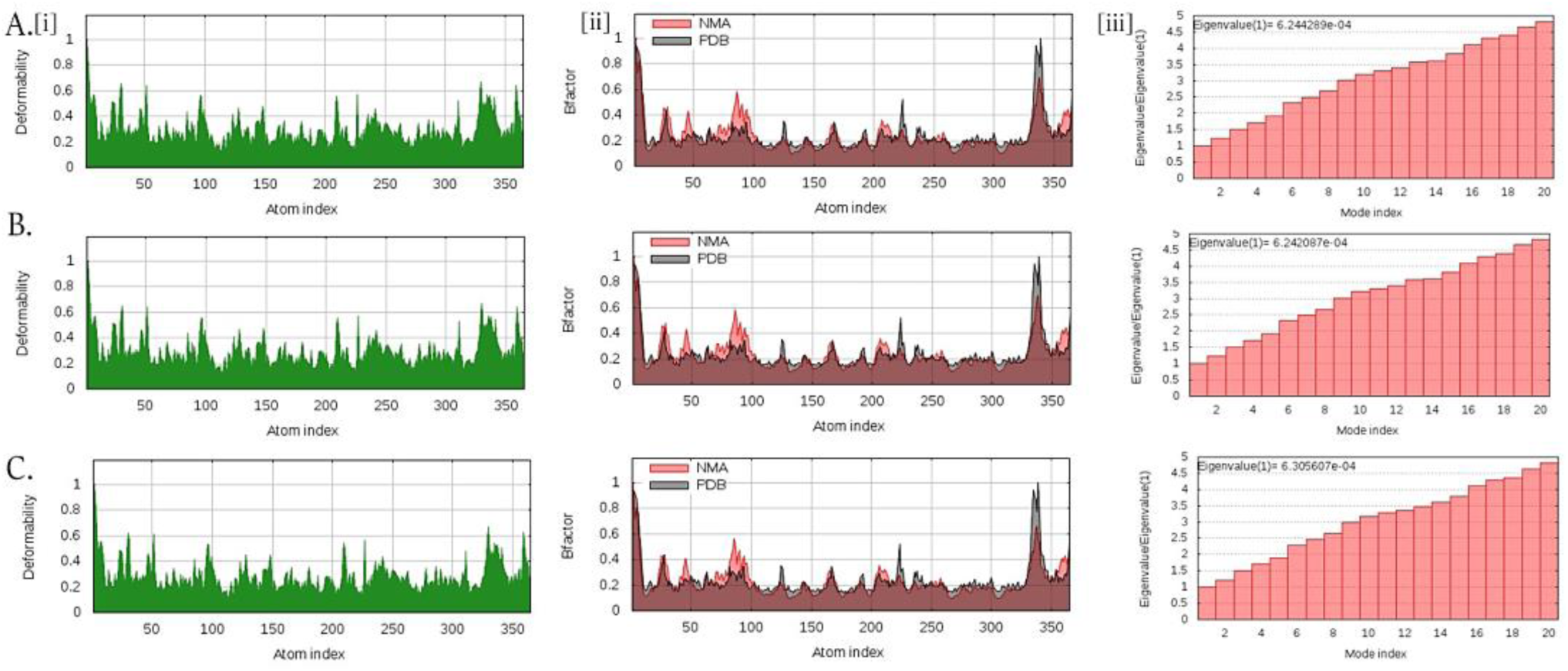
NMA analysis of a) 10651887 b) 14660440 C) CAY10603: i) deformability ii) B-factor iii) eigenvalues

The molecular dynamic simulation (NMA) of the complexes showed compound 14660440 to have an eigenvalue of 6.244289e-04, compound 10651887: 6.242087e-04, and compound CAY10603: 6.305607e-04. All the complexes showed similar eigenvalues however, compound 14660440 and 10651887 showed lower values.

### 3.4. Lead optimization

Bioisosteres are chemical moieties that are used to replace functional groups in compounds while retaining biological activity; bioisostere replacement is an established protocol for lead generation and optimization.^[58][59]^ Since benzimidazole has been suggested as a potential zinc-binding group^[60]^ and hydropyrimidine as an established capping group,^[26]^ we, therefore, held both regions immutable during the bioisostere replacement analysis; we sought to see if the binding affinity/binding affinity and specificity of the lead compounds would improve by substituting or modifying the linkers.

A total of 24 bioisostere replacement for compounds 1466044 were generated, while no suitable replacement was found for compound 10651887. These newly generated compounds were subjected to Molecular docking and QM-polarized docking, corresponding binding affinity was calculated. Of the 24 compounds generated 7 had QPLD binding affinity ranging between −81 to −70kcal/mol (Table 8). The bioisostere replacement protocol generally modified the amide linker of compound 1466044 with pyridazine and imine functional groups. Compounds 14660440_21 which had the highest QPLD binding affinity of −81.42kcal/mol (Table 8) had its amide linker replaced with pyridazine functional group which resulted in the formation of a π-π bond with LEY712 (Figure 11, 12), while compound 14660440_18 had its amide linker modified to an imine functional group forming carboximidamide which resulted in the formation of pi-cation interaction with PHE 643 and H-bound with SER 531 (Figure 11, 12). These replacements increased the QPLD binding affinity of un-optimized 14660440 from −70.80 to −81.42kcal/mol (Table 8).

**Table 8:** Compound 14660440 Bioisostere replaced compounds: docking score, binding affinity, and electronic properties.

| Compound ID | XP-docking Score (kcal/mol) | $\Delta G_{XP}$ (kcal/mol) | QPLD-docking score (kcal/mol) | $\Delta G_{QPLD}$ (kcal/mol) |
| --- | --- | --- | --- | --- |
| CAY10603 (Standard) | -8.494 | -90.70 | -8.810 | -98.55 |
| 14660440_21 | -10.445 | -85.45 | -10.281 | -81.42 |
| 14660440_18 | -10.346 | -80.43 | -9.827 | -81.37 |
| 14660440_13 | -9.992 | -80.35 | -10.211 | -67.92 |
| 14660440_19 | -9.236 | -79.74 | -9.434 | -51.19 |
| 14660440_24 | -9.612 | -79.55 | -10.043 | -73.46 |
| 14660440_3 | -10.935 | -78.84 | -9.866 | -67.85 |
| 14660440_14 | -9.77 | -78.46 | -9.835 | -71.39 |
| 14660440_23 | -9.574 | -78.43 | -9.382 | -78.96 |
| 14660440_16 | -10.018 | -78.32 | -9.305 | -68.45 |
| 14660440_5 | -9.933 | -78.19 | -9.181 | -71.42 |
| 14660440_11 | -9.827 | -78.12 | -9.045 | -70.98 |

**Figure 11:**
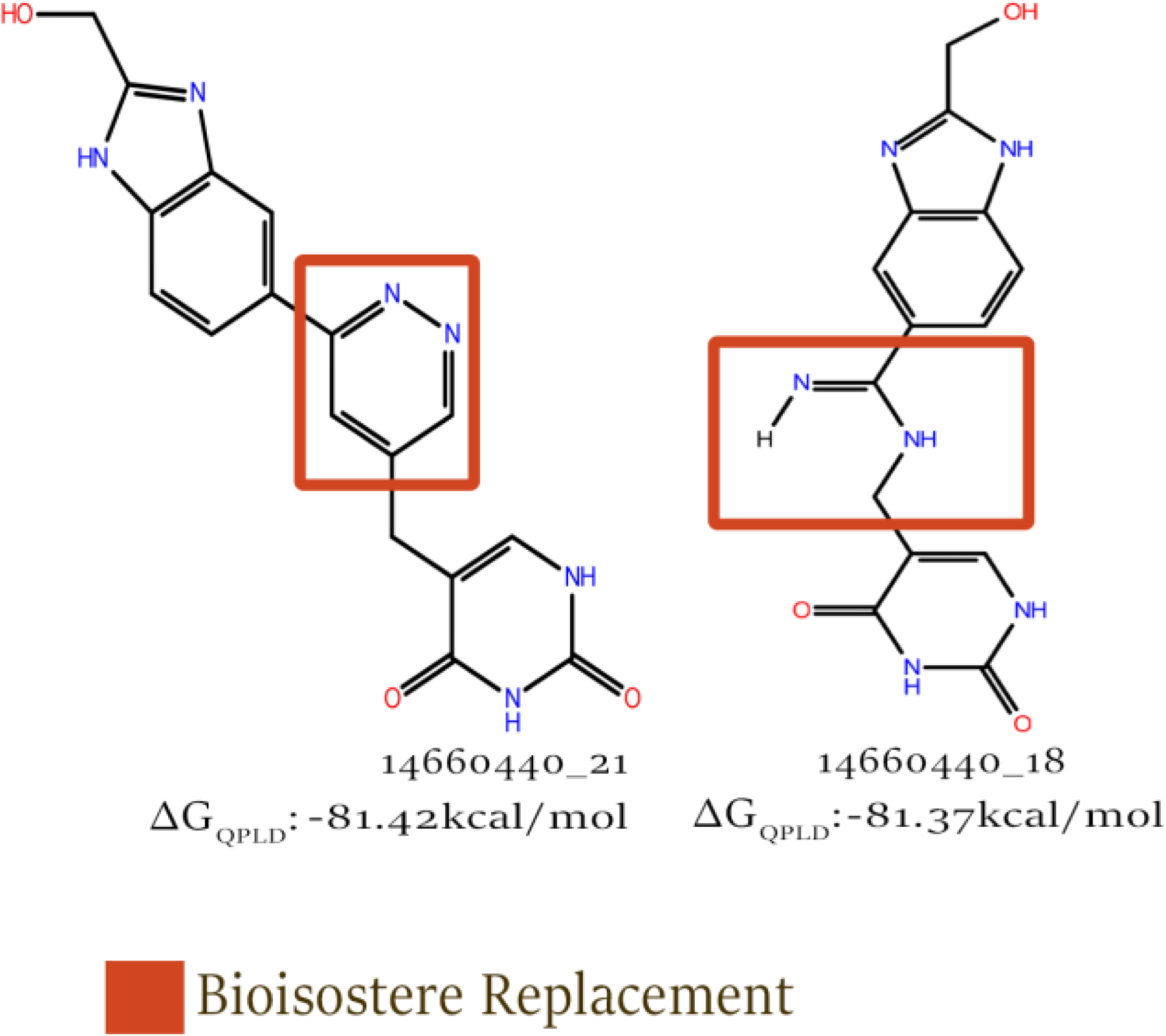
Top optimized compounds: 14660440_21 and 14660440_18.

**Figure 12:**
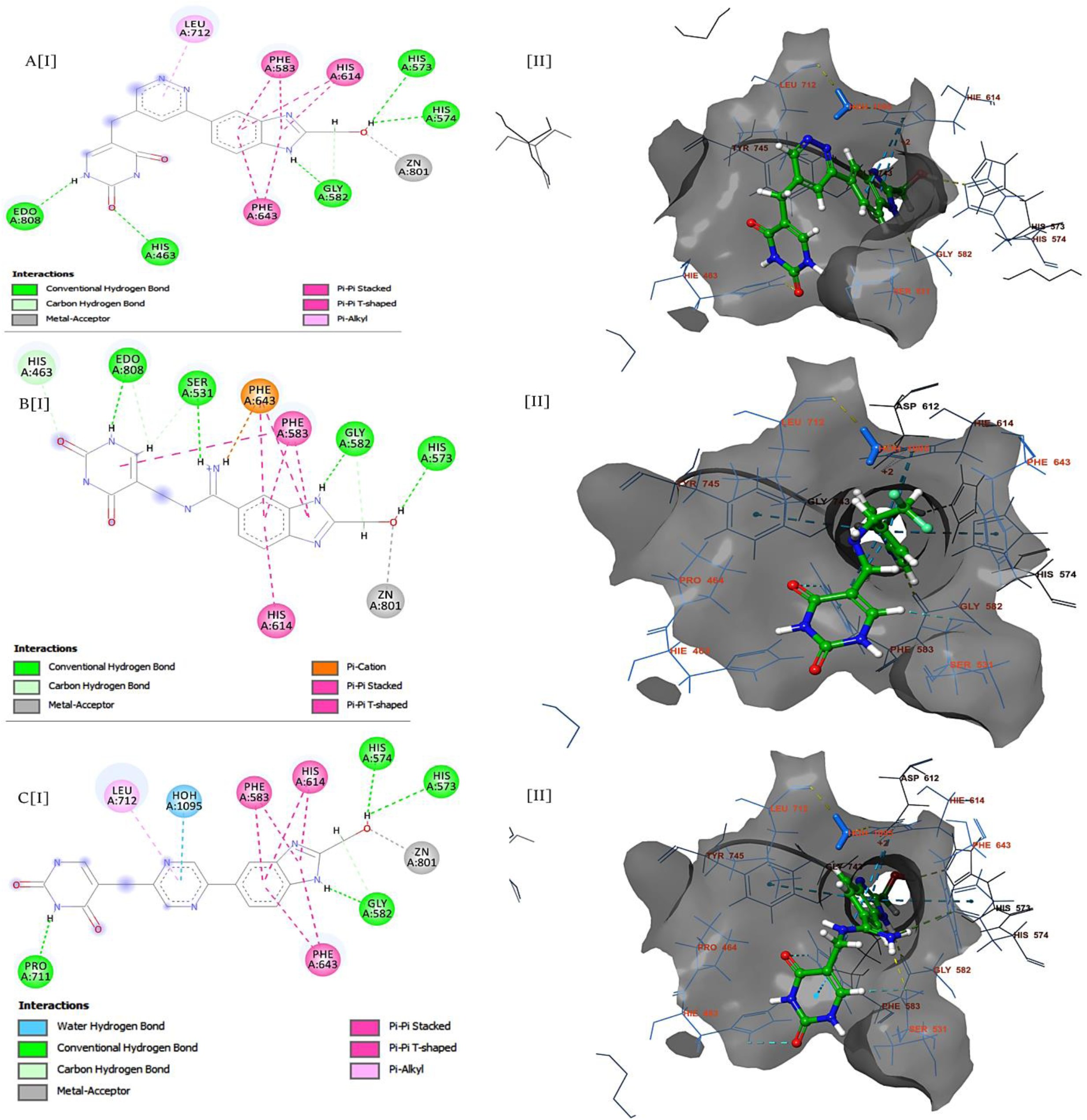
Compound 14660440 Bioisostere replacement compounds.

The specificity for HDAC6 also improved; the pyridazine linker replacement in 14660440_21 resulted in the formation of bonds with residue-specific to HDAC6: pi-alkyl bond was formed with LEU712 and H-bond was formed with HIS463 (Figure 12). The amine ion linker in 14660440_18 resulted in the formation of H-bond with HIS463 and SER531 (Figure 12).

### 3.5. Electronic Properties

Determining molecular orbitals and their properties help in understanding chemical reactive of organic compounds; of these orbitals, Highest Occupied Molecular orbitals (HOMO) and Lowest Unoccupied Molecular Orbital (LUMO) are the most important.^[61]^ The HOMO are low energy regions of a compound that readily donates electrons (nucleophile) while LUMO are high energy regions in a compound which are deficient in electrons, hence, readily accept electrons (electrophile); the LUMO is, therefore, the most reactive part of the compound.

The HOMO and LUMO of compounds 14660440_21 and 14660440_18 are visualized in figure 13. The figure shows HOMO of 14660440_21 to be around atoms of the pyridazine linker and LUMO to around atoms of the hydropyrimidine cap. HOMO of 14660440_18 is present around the benzimidazole zinc-binding pharmacophore; the LUMO is present around the imine linker. This, therefore, suggest the most reactive moiety of 14660440_21 is the hydropyrimidine cap while 14660440_18 is the imine linker (Figure 13).

**Figure 13:**
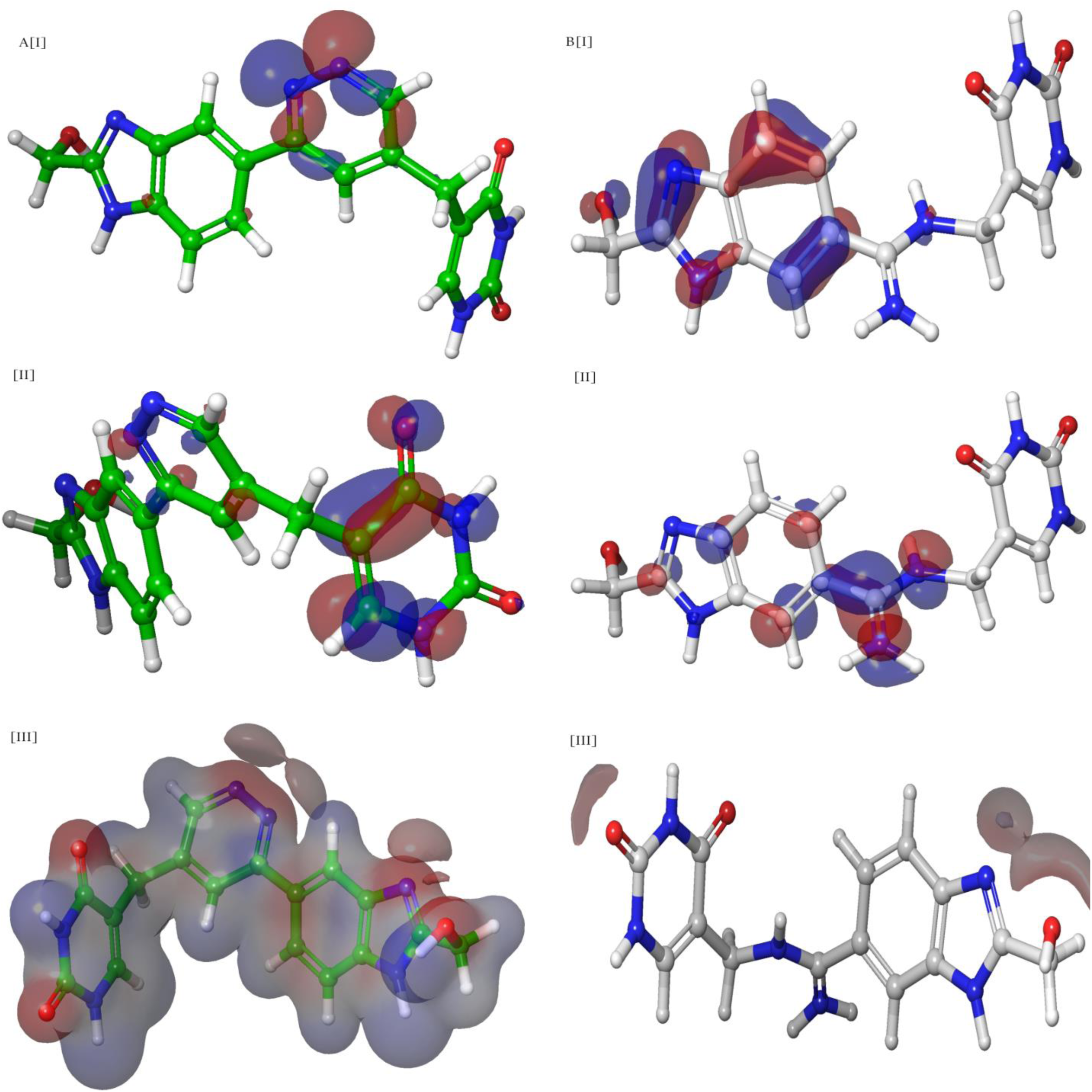
Electronic Properties of Compound a) 14660440_21 b) 14660440_18: i) HOMO ii) LUMO iii) MESP

From the calculated HOMO and LUMO energies different descriptors can be extrapolated; the HOMO-LUMO gap represents the difference between LUMO and HOMO energies, the gap indicates the ease by which electrons would move from HOMO to LUMO. A smaller gap indicates an easy movement of electrons which in turn indicates higher reactivity of the compounds. For both compound 14660440_21 and 14660440_18 HOMO-LUMO gap was 0.16 (Table 9).

**Table 9:** Electronic Descriptors of optimized 14660440 compounds.

| Compound ID | $\Delta G_{\text{QPLD}}$<br>(kcal/mol) | HOMO Energy<br>(eV) | LUMO Energy<br>(eV) | HOMO-LUMO<br>GAP | Ionization<br>energy<br>(I) | Electron<br>affinity<br>(A) | Chemical<br>potential<br>($\mu$) | Global<br>Hardness ($\eta$) | Global<br>Electrop<br>hilicity<br>($\omega$) |
| --- | --- | --- | --- | --- | --- | --- | --- | --- | --- |
| 14660440_21 | -81.42 | -0.22 | -0.06 | 0.16 | 0.22 | 0.06 | -0.14 | 0.08 | 0.12 |
| 14660440_18 | -81.37 | -0.35 | -0.19 | 0.16 | 0.35 | 0.19 | -0.27 | 0.08 | 0.25 |
| 14660440_13 | -67.92 | -0.23 | -0.04 | 0.19 | 0.23 | 0.04 | -0.14 | 0.10 | 0.10 |
| 14660440_19 | -51.19 | -0.14 | -0.09 | 0.05 | 0.14 | 0.09 | -0.12 | 0.03 | 0.24 |
| 14660440_24 | -73.46 | -0.23 | -0.06 | 0.17 | 0.23 | 0.06 | -0.15 | 0.09 | 0.13 |
| 14660440_3 | -67.85 | -0.34 | -0.18 | 0.16 | 0.34 | 0.18 | -0.18 | 0.08 | 0.20 |
| 14660440_14 | -71.39 | -0.22 | -0.04 | 0.18 | 0.22 | 0.04 | -0.13 | 0.09 | 0.09 |
| 14660440_23 | -78.96 | -0.22 | -0.05 | 0.17 | 0.22 | 0.05 | -0.14 | 0.09 | 0.11 |
| 14660440_16 | -68.45 | -0.21 | -0.04 | 0.17 | 0.21 | 0.04 | -0.13 | 0.09 | 0.09 |
| 14660440_5 | -71.42 | -0.23 | -0.05 | 0.18 | 0.23 | 0.05 | -0.14 | 0.09 | 0.11 |
| 14660440_11 | -70.98 | -0.21 | -0.21 | 0 | 0.21 | 0.21 | -0.21 | 0 | 0 |

Other descriptors that can be extrapolated from HOMO and LUMO energies are ionization energy (I), electron affinity (A), global hardness (η), chemical potential (µ), and global electrophilicity power of a ligand (ω) (see methods).^[62]^ Extremely hard compounds are weak electron acceptors, while a compound with lower chemical potential indicates good electrophiles the global electrophilicity power descriptor indicates the general reactivity of compounds.^[62]^ Compound 14660440_18 had the highest global electrophilicity power of 0.12, electron affinity: 0.19, and chemical potential: −0.27 (Table 9) making it the most reactive compound.

#### 3.5.1. Molecular electrostatic potential

Electrostatic potential explains the work done in moving a unit positive charge from infinity to a point, thus molecular electrostatic potential (MESP) is defined as the energy interaction between a positive charge and a molecule.^[62]^ MESP descriptor helps to identify regions in molecules that may be involved in electrophilic or nucleophilic attack thereby providing insights into reactivity of an organic compound.^[63]^ MESP provides a visual representation of positive and negative charge distribution in a molecule. Figure 13(a[iii], b[iii]) show MESP distribution of 14660440_21 and 14660440_18 respectively (Red: negatively charged, White: neutral, Blue: positively). Compound 14660440_21 was generally positive, with negative charges distributed over oxygen atoms on the hydropyrimidine cap, hydroxymethyl R group, and nitrogen atoms on benzimidazole (Figure 13a[iii]). Compound 14660440_18 however was predicted a cation with positive charges present over oxygen atoms on the hydropyrimidine cap and hydroxymethyl group on benzimidazole (Figure 13b[iii]).

### 3.6. ADMET Predictions

Having predicted the binding conformation, affinity, and mechanics of chemical reactivity, ADMET properties of the compounds were determined Using the discovery studio ADEMT descriptors calculator module (see methods). Lipinski drug-likeness and oprea lead-likeness descriptors were calculated using the MOE descriptor calculator (see methods).

All compounds except compound CAY10603 (Standard) passed the Lipinski drug-likeness test.^[63]^ CAY10603 and 14660440_18 failed oprea leadlike test ^[64]^ (Table 10). Aqoues absorption prediction, predicts the solubility of the compounds in water, which in turn determine oral absorption. The aqueous absorption ranged from good to optimal (Table 10). Intestinal absorption predictions predict the concentration that would be absorbed into the bloodstream. The predictions showed that intestinal absorption of CAY10603, 14660440, 14660440_5, and 14660440_18 might low (Figure 14; Table 10). Only compound 14660440_14 was predicted to be able to penetrate the blood-brain barrier (Figure 14). All other compounds (except CAY10603, 14660440, 14660440_5, and 14660440_18) where within 99% confidence ellipses of intestinal absorption i.e. 99% intestinal absorption is likely (Figure 14).

**Table 10:** ADMET Descriptor predictions.

| **Compound ID** | **$\Delta G_{QPLD}$** | **Lipinski**<br>**Druglike** | **Mutagenic** | **Oprea**<br>**Leadlike** | **Aqueous**<br>**Solubility** | **Cytochrome**<br>**P450 2D6**<br>**(CYP2D6)**<br>**Inhibition** | **Plasma**<br>**Protein**<br>**Binding** | **Human**<br>**Intestinal**<br>**Absorption** |
| --- | --- | --- | --- | --- | --- | --- | --- | --- |
| **CAY10603** | -90.6993 | True | True | False | Good | False | True | Low |
| **14660440** | -70.801 | True | False | True | Optimal | False | False | Low |
| **10651887** | -75.832 | True | False | True | Optimal | False | False | High |
| **14660440_21** | -81.4237 | True | False | True | Good | False | False | Moderate |
| **14660440_13** | -67.923 | True | False | True | Good | False | False | High |
| **14660440_18** | -81.3747 | True | False | False | Optimal | False | False | Low |
| **14660440_24** | -73.4641 | True | False | True | Good | False | False | Moderate |
| **14660440_16** | -78.5321 | True | False | True | Good | False | False | Moderate |
| **14660440_3** | -67.8505 | True | False | True | Good | False | False | High |
| **14660440_14** | -71.3854 | True | False | True | Good | False | False | High |
| **14660440_19** | -51.1944 | True | False | True | Optimal | False | False | Moderate |
| **14660440_23** | -78.963 | True | False | True | Good | False | False | Moderate |
| **14660440_16** | -68.4506 | True | False | True | Good | False | False | Moderate |
| **14660440_5** | -71.4231 | True | False | True | Good | False | False | Low |

**Figure 14:**
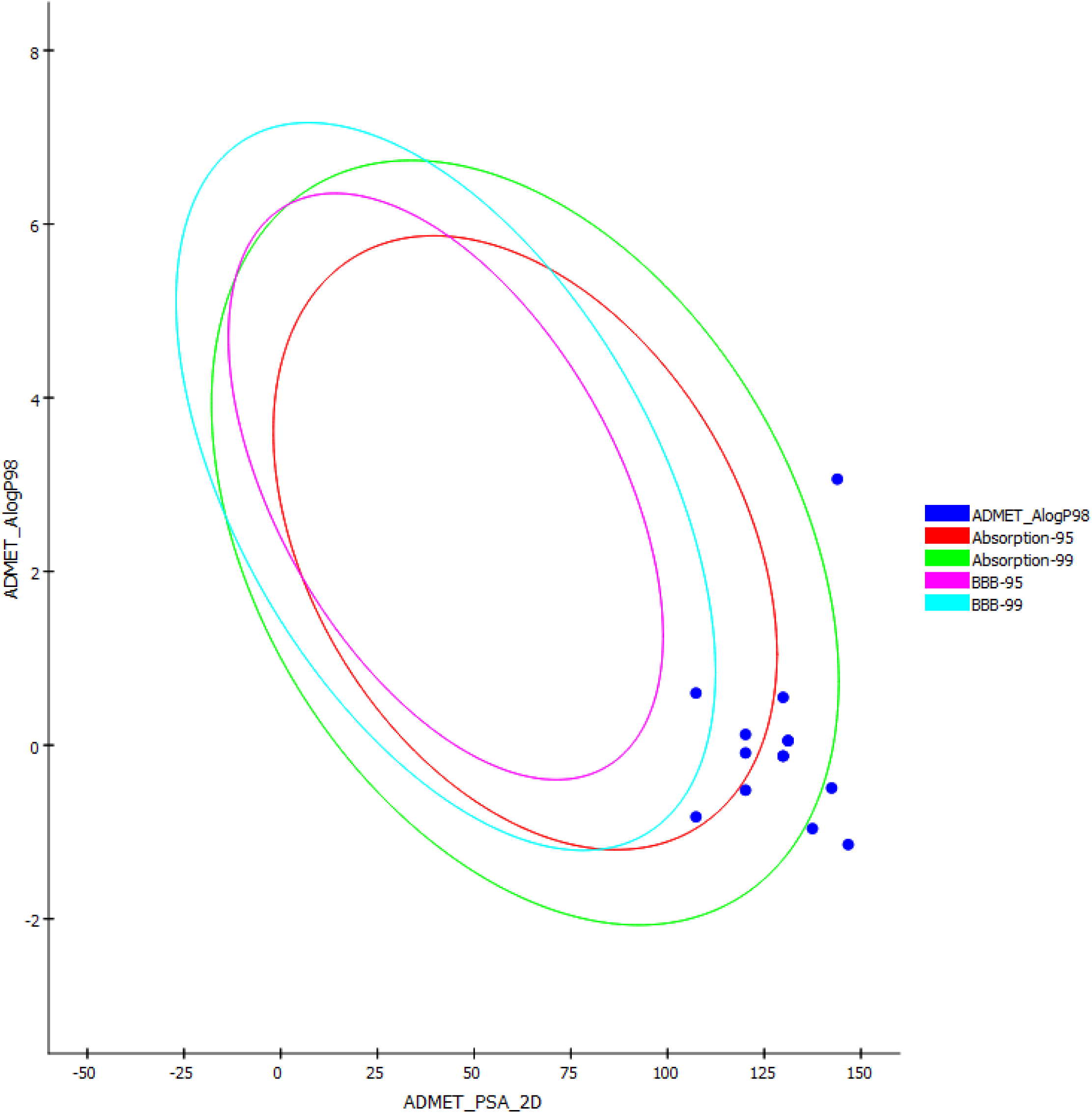
Human intestinal absorption and Blood-brain barrier penetration prediction plot: The ellipse describes the different regions in which well-absorbed compounds are expected. 95% of well-absorbed compounds are expected in the 95% ellipse and 99% of well-absorbed compounds are expected in the 99% ellipse.

Inhibition of cytochrome P450 2D6 (CYP2D6) by moieties of drugs is one of the major causes of drug-drug interaction^[65]^. Therefore, ensuring that this does not occur is an important aspect of drug optimization and design. None of the compounds were predicted to inhibit CYP2D6 (Table 10). Plasma protein binding (PPB) of the drug were also predicted. Plasma protein binding prediction is of utmost importance as the binding of compounds to plasma protein shields the compounds from metabolism and reduces their pharmacological efficacy.^[66]^ The prediction predicts whether 90% (and above) concentration of the compounds would exhibit PPB. Only Compound CAY10603 was predicted to exhibit plasma protein binding (Table 10). Mutagenicity of the compounds was also predicted. Only compound CAY10603 was predicted to be mutagenic (Table 10).

## 4.0. Discussion

Developing specific inhibitors for HDAC6 is still a daunting task due to the highly conserved nature of HDAC family protein. Based on literatures,^[26][67]^ the formation of non-covalent bonds with the following residues is key in discussing the putative mechanism of inhibition and specificity for compounds in this study:

- Catalytic residues: TYR 745; HIS 573; HIS 574; ZN^2+^; ASP 612; HIS 614; Catalytic H_2_O.
- Specificity residues: SER 531; HIS 462; HIS 463; PRO 464; LEU 712.

In this study, an RFC model was built on all know HDAC6 inhibitors (3742) deposited in Chembl (2003 to date). The model was validated on an external dataset using a 10 fold cross-validation, it had a balanced accuracy of 0.78, precision 0.79, recall 0.90, and AUC score of 0.768. The model was used to predict active compounds in the M sample representation of SCUBIDDOO; the predicted active compounds were docked and two compounds showing high docking score and fragment of the compounds having chelating interaction with ZN^2+^ were identified and used to search the SCUBIDOO database. A total of 7785 compounds having this fragment were downloaded and virtually screened, 24 compounds resulted and compound 14660440 and 10651887 was selected because they had the highest docking binding affinity of − 78.56kcal/mol and −78.21kcal respectively.

The two compounds were similar (RSMD: 0.61) in structure differing majorly in their linker moiety. They possessed a benzimidazole moiety with a 2-hydroxymethyl R group as their zinc-binding group, a hydropyrimidine as its capping group, an amide linker (14660440), and an amino linker (10651887). Recently Leandro *et al* ^[60]^ (using Fragment-based drug discovery) suggested benzimidazole functional group as a potential zinc-binding group. With this as proof of concept, we investigated a potential binding hypothesis using Induced-fit (IFD), QM-polarized docking (QPLD), and QM/MM optimization. The hypothesis suggests a possible mechanism of inhibition and specificity of the compounds by identifying consistent interaction. The criteria for selection were these:

- A binding interaction must have been observed consistently in Molecular docking, QPLD, IFD, and QM-MM predicted binding pose;
- Any new interaction observed in the IFD docking pose and consistent in QM-MM optimized pose is also selected.

Based on these above criteria compound 10651887 with a binding affinity of −75.83kcal/mol (QPLD) might inhibit HDAC6 by interacting with: HIS 573, HIS574, TYR 745, and ZN^2+^, and show specificity by interacting with SER531 and PRO464. Compound 14660440 with a binding affinity of −70.80kcal/mol (QPLD), interactions with catalytic HIS573, HIS614, TYR745, H_2_O, and ZN^2+^ might be its mechanism of inhibition. However, compound 14660440 did not interact with specificity residues. Considering the lack of specificity of compound 14660440 and the low binding affinities when compared with the standard (QPLD: −98.55kcal/mol), we sought to optimize the compounds using bioisostere replacement. No suitable bioisostere was found for compound 10651887, however, 24 new compounds were generated for compound 14660440. After molecular docking and QPLD the top two compounds where 14660440_21 (QPLD: − 81.42kcal/mol) and 14660440_18 (QPLD: −81.37kcal/mol) this two compounds showed a fold increase in binding affinity when compared with their parent compound (14660440: − 79.80kcal/mol). Compounds 14660440_21 had its amide linker replaced with a pyridazine functional group, this resulted in the formation of a π-π bond with LEY712 (specificity residue), and H-bond interactions with HIS463 (specificity residue) via its capping group (hydropyrimidine). Compound 14660440_18 had its amide linker modified with an imine functional group forming carboximidamide which resulted in the formation of H-bond with SER 531, and H-bond with HIS463 via its capping group (hydropyrimidine). Using Bioisostere replacement protocol we were successfully able to optimize and generate compounds with improved binding affinity and specificity.

Electronic descriptors were calculated to investigate the mode of the chemical reaction (nucleophilic or electrophilic), stability, and reactivity of the optimized compounds. Both compound 14660440_21 and 14660440_18 were shown to be very reactive and relatively stable (HOMO-LUMO gap: 0.16); compound 14660440_18 was predicted to be the most reactive (highest global electrophilicity power of 0.12). Based on HOMO-LUMO analysis the hydropyrimidine cap of 14660440_21 was predicted to be likely involved in electrophilic substitution while for 14660440_18, the imine linker was predicted to be the electrophilic region. The MESP analysis showed the hydropyrimidine cap, hydroxymethyl R group, and nitrogen atoms on benzimidazole to positively charge, suggesting that they might undergo electrophilic reactions. Compound 14660440_18 was predicted to be a cation suggesting that mode of reaction is likely to be purely electrophilic. ADMET predictions showed compound 14660440 and 10651887, with the optimized compounds to pass the drug-likeness test with no predicted mutagenicity, drug-drug interactions, and protein plasma binding. Oral absorption was predicted to be good and moderate for all the compounds, however, compound 14660440, 14660440_18, and 14660440_5 showed low intestinal absorption. Of note is that CAY10603 (Standard) was predicted to be a mutagenic and failed opera lead likeness test.

Having considered the binding interactions, electronic descriptors, and ADMET properties we suggest the following compounds as putative isospecific inhibitors of HDAC6: 10651887, 14660440_21, and 14660440_18. We consider these results to be predictive hence there is a need for experimental validations.

## 5.0. Conclusion

In this study, we have been able to build a random forest classifier model based on 3,742 inhibitors of HDAC6 developed from 2003 to date. The model had a 78% balanced accuracy after a 10 fold cross-validation on an external dataset; the model was used to screen the SCUBIDOO database. Two compounds were identified after virtual screening and were optimized for improved binding affinity and specificity. Electronic descriptors determined the mechanism of chemical reactivity and stability of the optimized compounds. Finally, ADMET properties were calculated with most of the compounds showing better drug properties than the standard.

## Supplementary materials

All supplementary materials are available in MendelyData at DOI: 10.17632/775s3xrhrk.1

- Jupyter notebooks and Python scripts
- Supplementary Tables and Figures
- SMILES of lead compounds and Bioisostere replacement optimized compounds
- Complete Data (Raw and Processed)

## Acknowledgment

We acknowledge the expert opinion of Idris Oluwaseun Mukhtar, George Oche Ambrose, and Alakanse Suleiman Oluwaseun.

## Notes

https://data.mendeley.com/datasets/775s3xrhrk/1

### Competing Interest Statement

The authors have declared no competing interest.

